# Evolution of Honey Resistance in Experimental Populations of Bacteria Depends on the Type of Honey, and Has no Major Side Effects for Antibiotic Susceptibility

**DOI:** 10.1101/2020.10.13.337063

**Authors:** Anna M. Bischofberger, Katia R. Pfrunder Cardozo, Michael Baumgartner, Alex R. Hall

## Abstract

With rising antibiotic resistance, alternative treatments for communicable diseases are increasingly relevant. One possible alternative for some types of infections is honey, used in wound care since before 2000 BCE and more recently in licensed, medical-grade products. However, it is unclear whether medical application of honey results in the evolution of bacterial honey resistance, and whether this has collateral effects on other bacterial traits such as antibiotic resistance. Here, we used single-step screening assays and serial transfer at increasing concentrations to isolate honey-resistant mutants of *Escherichia coli*. We only detected bacteria with consistently increased resistance to the honey they evolved in with two of the four tested honey products, and the observed increases were small (maximum two-fold increase in IC_90_). Genomic sequencing and experiments with single-gene knockouts showed a key mechanism by which bacteria increased their honey resistance was by mutating genes involved in detoxifying methylglyoxal, which contributes to the antibacterial activity of *Leptospermum* honeys. Crucially, we found no evidence that honey adaptation conferred cross-resistance or collateral sensitivity against nine antibiotics from six different classes. These results reveal constraints on bacterial adaptation to different types of honey, improving our ability to predict downstream consequences of wider honey application in medicine.

## Introduction

Antimicrobial resistance is one of the biggest challenges facing global public health (WHO, 2018). To preserve the effectiveness of antibiotics and to treat infections caused by resistant bacteria, alternative approaches are required that can be used instead of antibiotics or after they have failed. One possible alternative currently being investigated for some applications is honey (Descottes, 2009; Knipping *et al*., 2012; Vandamme *et al*., 2013). Produced by the honey bee, *Apis mellifera*, honey has a long history in human medicine (Breasted, 1948) and has remained a staple treatment in traditional medicine. More recently, medically certified honeys and honey-containing products have been licensed in various part of the world, primarily for topical application, such as in wound healing (Molan and Betts, 2004; Cooke *et al*., 2015), and treatment of otorhinolaryngological diseases (Werner and Laccourreye, 2011). The idea is appealing: honey is generally cheap and non-harmful to patients (Dunford and Hanano, 2004; Knottenbelt, 2014), inhibits bacterial growth (Basualdo *et al*., 2007; Carter *et al*., 2016) and can promote wound closure and healing (Molan, 1999). If honey could be used instead of antibiotics for some applications, this could contribute to managing antibiotic resistance. However, several open questions remain about whether bacteria exposed to inhibitory concentrations of honey evolve resistance to it, which genes or pathways are involved, and whether any such evolutionary responses have downstream effects on other properties relevant for treatment, in particular antibiotic resistance.

Despite open questions about how bacteria evolve honey resistance, the physiological causes of honey’s antibacterial activity have been investigated in various species, including human pathogens such as *Staphylococcus aureus, Pseudomonas aeruginosa, Streptococcus mutans* and *Enterococcus faecium* (Kwakman *et al*., 2008, 2011; Badet and Quero, 2011; Camplin and Maddocks, 2014). Multiple mechanisms have been implicated, including high sugar/low water content, acidity, hydrogen peroxide and non-peroxide molecules such as methylglyoxal (White *et al*., 1963; Allen *et al*., 1991; Molan, 1992; Mavric *et al*., 2008), with floral source of the honey being a major determinant of its mechanism of action (Allen *et al*., 1991; Lu *et al*., 2013; Maddocks and Jenkins, 2013). At the phenotypic level, various effects of honey have been reported across different species, including changes in cell-wall integrity and cell shape (Henriques *et al*., 2011; Brudzynski and Sjaarda, 2014; Wasfi *et al*., 2016), quorum sensing (Truchado *et al*., 2009; Lee *et al*., 2011; Wang *et al*., 2012), iron acquisition (Kronda *et al*., 2013; Ankley *et al*., 2020) and biofilm formation (Merckoll *et al*., 2009; Badet and Quero, 2011; Halstead *et al*., 2016). Perhaps unsurprisingly given this complex picture of inhibitory mechanisms, research to date on the evolution of honey resistance in bacteria has reached no clear consensus. For example, Cooper *et al*. (2010) reported no stable increase in resistance after exposure of various species to medical-grade *Leptospermum* honey *in vitro*, while Camplin and Maddocks (2014) and Lu *et al* (2019) detected increased resistance in *P. aeruginosa* cells recovered from honey-exposed biofilms *in vitro*. Thus, it remains unclear how rapidly bacteria exposed to honey can evolve reduced susceptibility to it. Another important question is whether adaptation to honey has side effects for antibiotic susceptibility. This is important for assessing the risk that honey application could contribute to the spread of antibiotic resistance, affecting the success of antibiotic treatments used in combination or later against the same bacteria. There is some evidence that honey resistance affects susceptibility to rifampicin and imipenem in *P. aeruginosa* (Camplin and Maddocks, 2014). To understand the general picture of how honey resistance affects antibiotic susceptibility, we need to test a wider range of bacteria and antibiotics, and characterise the genetic pathways by which bacteria become resistant.

We chose to study *E. coli* because it is a common pathogen in humans and animals (Dowd *et al*., 2008; Suojala *et al*., 2013), frequently associated with surface wounds and infections of the intestinal tract that might be suitable for honey treatment (Haffejee and Moosa, 1985; Willix *et al*., 1992; Carnwath *et al*., 2014). Antibacterial resistance has increased in *E. coli* (Tadesse *et al*., 2012; O’Neill, 2014), and many of the resistance mechanisms found in *E. coli* can also be found in other species (Philippon *et al*., 1989; Paulsen *et al*., 1996), suggesting it as a good model for studying resistance evolution. To find out how bacteria respond to honey exposure phenotypically and genotypically, we experimentally evolved *E. coli* in the presence of four different honeys (two medical-grade honeys, two commercially available honeys) by gradually exposing bacteria to increased honey concentrations during serial passage. We then measured honey susceptibility of evolved bacteria from this experiment, as well as population growth in the absence of honey. To identify genes involved in honey resistance, we used whole-genome sequencing of these evolved isolates, as well as single-gene knockout variants. We also used a second screen for honey-resistant mutants, by exposing many replicate populations to high honey concentrations in a single-step (selective plating of overnight cultures). Lastly, we tested for collateral effects of honey adaptation on antibiotic resistance by determining the phenotypic resistance of honey-adapted isolates to antibiotics of different classes. Our results show that, even upon gradually increasing exposure, large changes in honey resistance in *E. coli* populations growing *in vitro* are rare. However, for some honey products we identified mechanisms driving moderate increases in honey resistance, and we find no indication for cross-resistance between honey and antibiotics.

## Materials and Methods

### Organisms and Growth Conditions

We used *E. coli* K-12 MG1655 as parental strain for the evolution experiment and isolation of single-step mutants. We used *E. coli* K-12 BW25113 and single-gene knockout variants derived from it (Keio Knockout Collection (Baba *et al*., 2006)) for knockout experiments. We stored all isolates in 25% glycerol at −80°C. We performed routine culturing in lysogeny broth (LB, Sigma-Aldrich (Merck KGaA, Germany)) at 37°C with shaking at 180rpm.

### Honeys and Antibiotics

The different honey products we used are listed in Table 1. Honeys were stored in a cool, dark place and, in the case of medical-grade honeys, opened tubes were only used as long as recommended by the manufacturer (SurgihoneyRO™: 4 weeks; Medihoney™ Medical Honey: 4 months). Because the best-before date on commercial honeys does not concern their antimicrobial activity but its edibility, and because prolonged storage can affect hydrogen peroxide content of honey (Irish *et al*., 2011), non-medical-grade honeys were also used for a maximum of four months after opening. After plating honey samples on agar plates (LB broth with agar (Lennox) (Sigma-Aldrich (Merck KGaA, Germany)) at 35g/L), we observed colony-forming units with the commercial honey and Manuka honey. This was no longer the case after filtering honey solutions with a Filtropur S 0.45 filter (Sarstedt, Germany) (Wasfi *et al*., 2016). Accordingly, for all four honeys, honey-containing growth media were prepared immediately before the start of each experiment by diluting honey in LB (pre-heated to 55°C) and filter sterilizing. We purchased amoxicillin (product number A8523), chloramphenicol (product number 23275), ciprofloxacin (product number 17850), gentamicin (product number 48760), kanamycin (product number 60615), neomycin trisulfate salt hydrate (product number N1876), polymyxin B (product number 5291), tobramycin (product number T4014), and trimethoprim (product number 92131) from Sigma-Aldrich (Merck KGaA, Germany). We prepared antibiotic stock solutions at the outset of the experiments and filter sterilized (Filtropur S 0.2 (Sarstedt, Germany)) and stored them according to the manufacturers’ instructions (stock solutions: amoxicillin 25mg/mL in sterile distilled water (dH_2_O); chloramphenicol 50mg/mL in 70% ethanol; ciprofloxacin 20mg/mL in 0.1M HCl; gentamicin 50mg/mL in dH_2_O; kanamycin 40mg/mL in dH_2_O; neomycin 40mg/mL in dH_2_O; polymyxin B 20mg/mL in dH_2_O; tobramycin 40mg/mL in dH_2_O; trimethoprim 25mg/mL in DMSO).

**Table 1.** Honey products used in this study.

| Product name | Product description | Initial IC<sub>90</sub> for *E. coli* K-12 MG1655 | Manufacturer | Purchased from |
| --- | --- | --- | --- | --- |
| *SurgihoneyRO*<sup>TM</sup> | medical honey; sterile, bio-engineered; undiluted; single-origin (unspecified) | 5%(w/v) | Maotoke Holdings Ltd, Abingdon, United Kingdom | tubes of 20g and 50g: H&R Healthcare, Hull, United Kingdom |
| *Medihoney*<sup>TM</sup> Antibacterial Medical Honey | medical honey; sterile, undiluted; *Leptospermum* honey (methylglyoxal content unspecified) | 16%(w/v) | sorbion austria, Zwölfaxing, Austria | tubes of 20g: Puras AG, Bern, Switzerland; tubes of 50g: The Honeydoctor, The Littledart Company Ltd, Tiverton, United Kingdom, and puravita ag, Speicher, Switzerland |
| *MGO*<sup>TM</sup> Manuka Honey *MGO*<sup>TM</sup>550+ | consumer product; natural, undiluted; *Leptospermum* honey (methylglyoxal content of 550 mg/kg) | 10%(w/v) | Manuka Health New Zealand Ltd, Te Awamutu, New Zealand | 250g jars: nu3 GmbH, Berlin, Germany |
| Commercial honey (Fairtrade Liquid Blossom Honey) | consumer product; natural, undiluted; poly-origin (Chili, Guatemala, (Mexico)) | 10%(w/v) | Coop, Basel, Switzerland | 550g jars: Coop, Basel, Switzerland |

### Measuring Susceptibility of Isolates to Different Honeys

We used the 90% inhibitory concentration (IC_90_) of each antibacterial compound as an indicator of resistance. We defined the IC_90_ as the lowest concentration tested above which bacterial growth did not exceed 10% of growth of the same isolate in the absence of antibacterials (i.e. none of the tested concentrations at or above the IC_90_ supported >10% growth; this definition is used in the results sections below). A minority of dose-response curves were not monotonic, in that some individual concentrations below the IC_90_ inferred using the above definition supported <10% growth; this did not affect our overall conclusions (checked by using an alternative definition of the IC_90_ as the lowest individual concentration supporting <10% growth, which supported the same conclusions). We estimated the IC_90_ towards four honey products for ancestral strain *E. coli* K-12 MG1655 (assay A); for 18 single-step putative resistant mutants (assay B, also including the ancestral strain); for 14 serially-passaged putative honey-resistant mutants and six serially -passaged control isolates (assay C, also including the ancestral strain); and for 28 single-gene knockout variants and their ancestral strain *E. coli* BW25113 (assay D, for details on selection of single-gene knockout variants see below) by measuring their growth in liquid culture at different concentrations. For each assay, we transferred independent LB-overnight cultures (cultured in flat-based 96-well microplates (Sarstedt, Germany)) into microplates filled with various honey concentrations and plain LB as a control, using a pin replicator (1/100 dilution, 2μL in 200μL). We used slightly different concentration ranges in different sets of assays (range of tested concentrations is given in Table S1). The assays were conducted with independent controls (ancestral strain) on different days. After inoculating assay microplates, we incubated them overnight at 37°C and quantified bacterial growth by measuring optical density at 600nm (OD_600_) with a microplate reader (Infinite^®^ 200 PRO, Tecan Trading AG, Switzerland) at the beginning and end of the experiment (0h and 24h). We corrected OD_600_ scores for the optical density of the media. In each assay, we assessed multiple replicates of each strain-compound combination (<u>assay A</u>: three replicates; <u>assay B</u>: five replicates; <u>assay C</u>: four replicates; <u>assay D</u>: three replicates).

### Experimental Evolution (Fig. S1A)

We serially passaged multiple selection lines of *E. coli* K-12 MG1655 in filtered solutions of each of the four honey products and in the absence of honey for 22 days, transferring daily (4 honey treatments + 1 control treatment = 5 evolution environments). In summary: at each transfer, each selection line was inoculated into multiple wells containing various honey concentrations. After overnight incubation, we transferred from the well with the highest concentration supporting viable growth (Fig. S1A). In more detail: to begin the experiment, we streaked out *E. coli* K-12 MG1655 from glycerol stocks onto LB agar plates. After overnight incubation, we inoculated six selection lines in each evolution environment, each with an independent colony (5 evolution environments x 6 colonies = 30 selection lines). In this first step, we cultured each selection line for 2h in 5mL of LB at 37°C with shaking at 180rpm. Then, for every selection line, we inoculated seven microplate wells filled with 200μL LB. After overnight incubation, we transferred the seven cultures of each selection line (5μL of each culture) into a fresh microplate filled with: 200μL of unsupplemented LB (“rescue well”), LB supplemented with one of five concentrations of the respective honey product, or honey stock solution (30 or 50 %(w/v)). We incubated microplates overnight at 37°C and quantified bacterial growth by measuring OD_600_ after 0h and 24h. On the following days, for every selection line, we determined the well at the highest honey concentration where ΔOD_600_ (OD_600_ 24h – OD_600_ 0h) > 0.1 (an arbitrary cut-off we took as an indication of viable growth) and transferred 5μL to seven wells in a new microplate. In cases where ΔOD_600_ in all honey-supplemented wells was < 0.1, we used the rescue-well culture to inoculate the fresh microplate. To gradually expose selection lines to higher concentrations of honey, we adjusted the range of concentrations tested over time, according to the performance of individual selection lines (concentrations and OD scores over time are given in Fig. S2). In the control treatment (no honey), we used a single well for each selection line at each transfer. We did this for 22 days, freezing microplates from days 3, 6, 9, 12, 15, 18, 21 and 22.

At the end of the experiment, we isolated a single colony at random from each selection line. On day 22, we streaked out a sample from the well with the highest honey concentration where ΔOD_600_ > 0.1 onto LB agar. After overnight incubation, we picked one colony per selection line, grew it overnight in 5mL LB and stored it at −80°C. We used colony PCR (primer sequences: forward: 5’-AGA CGA CCA ATA GCC GCT TT-3’; reverse: 5’-TTG ATG TTC CGC TGA CGT CT-3’) to ensure that all colony isolates were *E. coli* K-12 MG1655. For five selection lines (Medihoney_01, Commercial_02, Commercial_03, Commercial_04, Commercial_05), we found no colonies when streaking out samples on LB agar at day 22. We initially screened for honey-resistant phenotypes of colony isolates from the other selection lines by culturing each colony isolate in honey-supplemented medium at a concentration in which the parental strain was not able to grow. This led us to exclude five isolates that did not grow at honey concentrations above that of the parental strain (Surgihoney_01, Surgihoney_02, Surgihoney_03, Surgihoney_06, Commercial_01). We then proceeded with the remaining 14 serially-passaged putative honey-resistant mutants, plus six control isolates serially passaged in LB, sequencing all 20 genomes and quantifying their honey susceptibility as described below/above respectively.

### Genetic Analysis of Serially-Passaged Mutants

We sequenced the 14 serially-passaged, putative resistant mutants and the six LB-adapted control isolates. Overnight cultures inoculated with single colonies were centrifuged at 5000xg at room temperature for 10 min. After removal of the supernatant, we stored cell pellets at −20°C until further processing. We used the QIAGEN Genomic-tip 20/G (Cat. No. 10223, Qiagen, the Netherlands) according to the manufacturer’s instructions for genomic DNA (gDNA) extraction. In brief: We resuspended the bacterial cell pellets in 1mL Buffer B1, 2μL RNase A solution (100mg/mL), 20μL lysozyme (100mg/mL) and 45μL Proteinase K (20mg/mL). Afterwards, we incubated them at 37°C for up to 1h. Then, we added 350μL of Buffer B2 and mixed thoroughly by inverting the tubes several times and vortexing them a few seconds. Following incubation at 50°C for up to 1h, we loaded the lysates onto the pre-equilibrated QIAGEN Genomic-tips and left the samples to pass the resin by gravity flow. We washed the QIAGEN Genomic-tips thrice with 1mL Buffer QC to remove any remaining contaminants. We eluted the DNA twice with 1mL Buffer QF pre-warmed to 50°C, discarded the Genomic-tips and precipitated the DNA by adding 1.4mL room-temperature isopropanol to the eluted DNA. We precipitated the DNA by inverting the tube 10-20 times and spooled the DNA using a glass rod. We immediately transferred the spooled DNA to a microcentrifuge tube containing 160μL elution buffer (Buffer EB: 10 mM Tris-Cl, pH 8.5) and dissolved the DNA overnight on a shaker (20rpm). We quantified the obtained gDNA using the Quant-iTTM dsDNA BR (Broad Range) Assay Kit (Thermo Fisher Scientific, USA) in the QubitTM Fluorometer (Thermo Fisher Scientific). We used a Nanodrop (Thermo Fisher Scientific) to control the purity of gDNA (ratios A_260_/A_280_ and A_260_/A_230_ ≥ 1.8). We sequenced at the Functional Genomic Center, Zurich, Switzerland, using the Illumina Hiseq 4000 platform after library preparation with the Nextera XT DNA Library Prep kit (Illumina, USA).

We trimmed and quality-filtered all sequences with trimmomatic (Bolger *et al*., 2014) with the following parameters: ILLUMINACLIP:<NexteraPE adapters fasta file>2:30:10; LEADING: 3; TRAILING: 3; SLIDINGWINDOW:4:15; MINLEN:80.. We mapped the reads of the ancestral strain of our resistance isolation experiment (2.5 x 10^6^ reads) against the reference sequence of *E. coli* K-12 MG1655 (NCBI accession number: U00096) using breseq 0.33.1 (Deatherage and Barrick, 2014). We used gdtools implemented in breseq to integrate the identified mutations into the reference genome. For variant calling, we mapped all reads of the serially-passaged and single-step putative resistant mutants (average number of reads per sample: 4.1 x 10^6^ ± 1.5 x 10^6^) against the refined reference genome using breseq. The sequencing data has been deposited in the European Nucleotide Archive under the study accession number PRJEB35347 (https://www.ebi.ac.uk/ena).

### Experiments with Single-Gene Knockout Variants

After identifying genes that potentially contribute to honey adaptation (Table S2), we tested for further evidence of the role of these genes in honey resistance using single-gene knockout variants from the Keio Knockout Collection (Baba *et al*., 2006) for *E. coli* K-12. We tested for a change in the resistance phenotype of these knockout variants relative to the ancestral strain of the knockout collection, *E. coli* K-12 BW25113, using the resistance phenotyping assay described above (assay D). When choosing genes to investigate, we concentrated on those (1) which were affected by independent mutations in at least two selection lines, (2) where mutations were not detected in isolates from the control treatment, (3) which were not annotated as “pseudogene”, “intergenic” nor “non-coding”, and (4) for which there is an available knockout variant in the Keio Collection (limited to non-essential genes).

### Measuring Population Growth of Serially-Passaged Mutants in the Absence of Honey

We tested whether our serially passaged isolates showed altered population growth in the absence of honey relative to the ancestral strain. To do this, we grew four independent cultures (each inoculated from a different colony) of each serially-passaged isolate (including control-evolved isolates, *n* = 20) and the ancestral isolate, each in 150μL LB in a microplate in a randomized layout. After overnight incubation, we used a pin replicator to inoculate a fresh microplate (all wells filled with 150μL LB). We then measured OD_600_ every 15min for 24h (shaking before each measurement).

### Measuring Susceptibility of Serially-Passaged Mutants to Antibiotics

We measured the phenotypic resistance (IC_90_) of the 20 serially-passaged isolates (14 putative resistant mutants, six LB-adapted control isolates) and of the parental strain (*E. coli* K-12 MG1655) for nine antibiotics representing six different classes: amoxicillin (penicillin), ciprofloxacin (fluoroquinolone), chloramphenicol (chloramphenicol), gentamicin (aminoglycoside), kanamycin (aminoglycoside), neomycin (aminoglycoside), polymyxin B (polymyxin), tobramycin (aminoglycoside), and trimethoprim (dihydrofolate reductase inhibitor). We decided to test several aminoglycoside drugs (gentamicin, kanamycin, neomycin, tobramycin) because previous studies have found that bacteria exposed to honey reduce the expression of genes involved in the TCA cycle (Lee *et al*., 2011; Jenkins *et al*., 2014), while others report a link between aminoglycoside susceptibility and defects or down-regulated gene expression in the bacterial respiratory chain, including the TCA cycle (Magnet and Blanchard, 2005; Chittezham Thomas *et al*., 2013; Shan *et al*., 2015; Su *et al*., 2018; Zhou *et al*., 2019). We conducted the assays using a similar protocol as described above for honey. In brief, we first incubated independent replicate populations of each isolate in a randomized layout in microplates overnight. From these microplates we inoculated assay plates filled with LB supplemented with antibiotics at various concentrations. With 2-fold broth dilution, the non-zero concentration ranges were: amoxicillin 128 – 4 μg/mL, chloramphenicol 32 – 1 μg/mL, ciprofloxacin 1 – 0.0.03125 μg/mL, gentamicin 32 – 1 μg/mL, kanamycin 32-1 μg/mL, neomycin 64 – 2 μg/mL, polymyxin B 4 – 0.125 μg/mL, tobramycin 32 – 1 μg/mL, trimethoprim 4 – 0.125 μg/mL. We measured bacterial growth by the change in OD_600_ (0h, 24h) as described above. We conducted the assays for all antibiotics on the same day.

### Single-Step Isolation of Honey-Resistant Mutants (Fig. S1B)

As a second screen for mutants of *E. coli* K12 MG1655 with increased honey-resistance, we plated aliquots of multiple independent overnight cultures, grown in the absence of antibiotics, on LB agar supplemented with each honey product.

For each honey type, we first grew 54 independent overnight cultures (250μL per culture in LB in a 96-well microplate; Fig. S1B). We then plated each culture onto a separate honey-supplemented LB agar plate (prepared in 6-well culture plates (Sarstedt, Germany)), plating 100μL of 18 separate cultures at each of three concentrations per honey product. We prepared honey-supplemented LB agar by adding LB-honey solution (at double the concentration of the desired final concentration in the plates, prepared as described above) to hand-warm double concentrated LB agar (i.e., at 70g/L agar). After mixing, we added 4mL of this honey-supplemented agar to wells. We used honey concentrations 1.25-, 1.5-, 2- or 3-times higher than previously determined IC_90_s of the wild-type strain in liquid. We incubated the plates overnight at 37°C before checking for resistant mutants. For SurgihoneyRO™, Medihoney™ and Manuka honey, we picked six putative honey-resistant colonies, each from a separate well. When isolating colonies, we prioritized those from wells with higher concentrations of honey. For commercial honey, we observed no viable colonies on honey-supplemented agar despite four attempts on different days (total of 216 cultures). We cultured the selected putative resistant colonies overnight in 5mL LB and suspended them in 25% glycerol for storage at −80°C. We then tested these 18 single-step putative honey-resistant mutants for phenotypic resistance as described above. We also used a second test to see if putative honey resistance phenotypes were robust, by streaking out frozen stocks of each colony isolate on honey-supplemented agar at a concentration inhibitory to the parental strain. Only three of 18 single-step putative honey-resistant mutants were able to form colonies under these conditions (Medihoney_10, Surgihoney_09, Surgihoney_10).

### Statistical Analysis

#### Phenotypic honey resistance of serially-passaged mutants

To test whether isolates from different evolution environments had different honey-resistance profiles, we used a linear-mixed effects model (lmer function in R’s lmerTest package (R version: 4.0; package version: 3.1-2)), with evolution environment and assay compound as fixed effects and genotype (isolate) as a random effect. We compared models with and without the interaction between fixed effects, using the anova function of the stats package to test significance. When looking at individual assay compounds, we used evolution environment as a fixed effect and genotype as a random effect. We used maximum likelihood estimation (*REML = F*). Wild-type data was excluded from this analysis; we tested for differences in IC_90_ between evolved isolates and the wild type separately, with *t*-tests (*p*-values adjusted for multiple testing using the Holm-Bonferroni method).

#### Population growth of serially-passaged mutants in absence of honey

We used R’s nls and SSlogis functions in the stats package to estimate the growth rate and yield for each culture, with yield corresponding to the *Asym* parameter of the models (= asymptote) and growth rate corresponding to the inverse of the *scal* parameter (= numeric scale parameter on input axis). We tested for a difference in growth parameters between each evolved isolate and the parental strain with a *t*-test, adjusting *p*-values using sequential Bonferroni correction. We also tested for an average difference in growth rate or yield among evolved isolates from different evolution environments. We did this using a linear mixed-effects model, with evolution environment as a fixed effect and genotype as random effect, and excluding data from the wild type. We used maximum likelihood estimation (*REML = F*) with the lmer function in the lmerTest package.

#### Phenotypic resistance profiles of serially-passaged mutants (antibiotics)

We conducted an Analysis of Variance (ANOVA), using the aov function in R’s stats package, to test whether there is cross-resistance between honey and antibiotics, with data on the phenotypic antibiotic resistance of isolates serially passaged in different honeys. We tested each evolved-isolate-versus-wild-type combination separately, with genotype (evolved vs wild type) and assay compound (antibiotic) as factors, including the interaction term and with *p*-values adjusted using the Holm-Bonferroni method.

## Results

### Experimental Evolution of Manuka Honey-Resistant Bacteria by Serial Passaging

During serial passage at gradually increasing honey concentrations (Fig. S1A), most selection lines showed improved population growth at concentrations that were initially inhibitory to the ancestral strain (Fig. S2 & S3). When we measured the phenotypic resistance of 14 putative honey-resistant mutants and six control isolates, each isolated from a different selection line after 22 days, several showed increased resistance compared to the ancestral strain against one or more honey products (Fig. 1 & S4). The average change in resistance varied among evolved isolates passaged with different honey compounds (effect of evolution environment: F_4, 20_ = 1.799, p > 0.1). These differences also depended on which honey compound was used in the assay (evolution environment × assay environment interaction: χ^2^(12) = 234.99, p < 0.001). We observed the largest change in resistance for mutants selected with Manuka honey or Medihoney™ when they were assayed with Manuka honey (mean change relative to the ancestor of 2.13-fold (s.d. = ±0.22) and 2.07-fold (s.d. = ±0.2), respectively). Manuka honey resistance of Manuka honey- and Medihoney™-evolved isolates was also significantly higher than for control-evolved isolates from the honey-free LB-medium treatment (t_9.9_ = 6.71, p < 0.001 and t_9_ = 6.15, p < 0.001, respectively). On average, Manuka honey-evolved and Medihoney™-evolved isolates also had moderately increased resistance to Medihoney™ (mean change relative to the ancestor of 1.15-fold (s.d. = ±0.16) and 1.17-fold (s.d. = ±0.1), respectively), but this was not significantly different from that of control-evolved isolates on average (Manuka: t_8.1_ = −0.24, p > 0.05; Medihoney™: t_8.2_ = 0, p > 0.05). Some other individual isolates had consistently increased Manuka honey or SurgihoneyRO™ resistance (all replicates for a given isolate higher than all replicates for the ancestor; Fig. 1 & S4), but these changes were small compared to those for Manuka honey- and Medihoney™-evolved isolates tested with Manuka honey. Thus, after experimental evolution we observed the strongest evidence of honey adaptation with Manuka honey.

**Figure 1.**
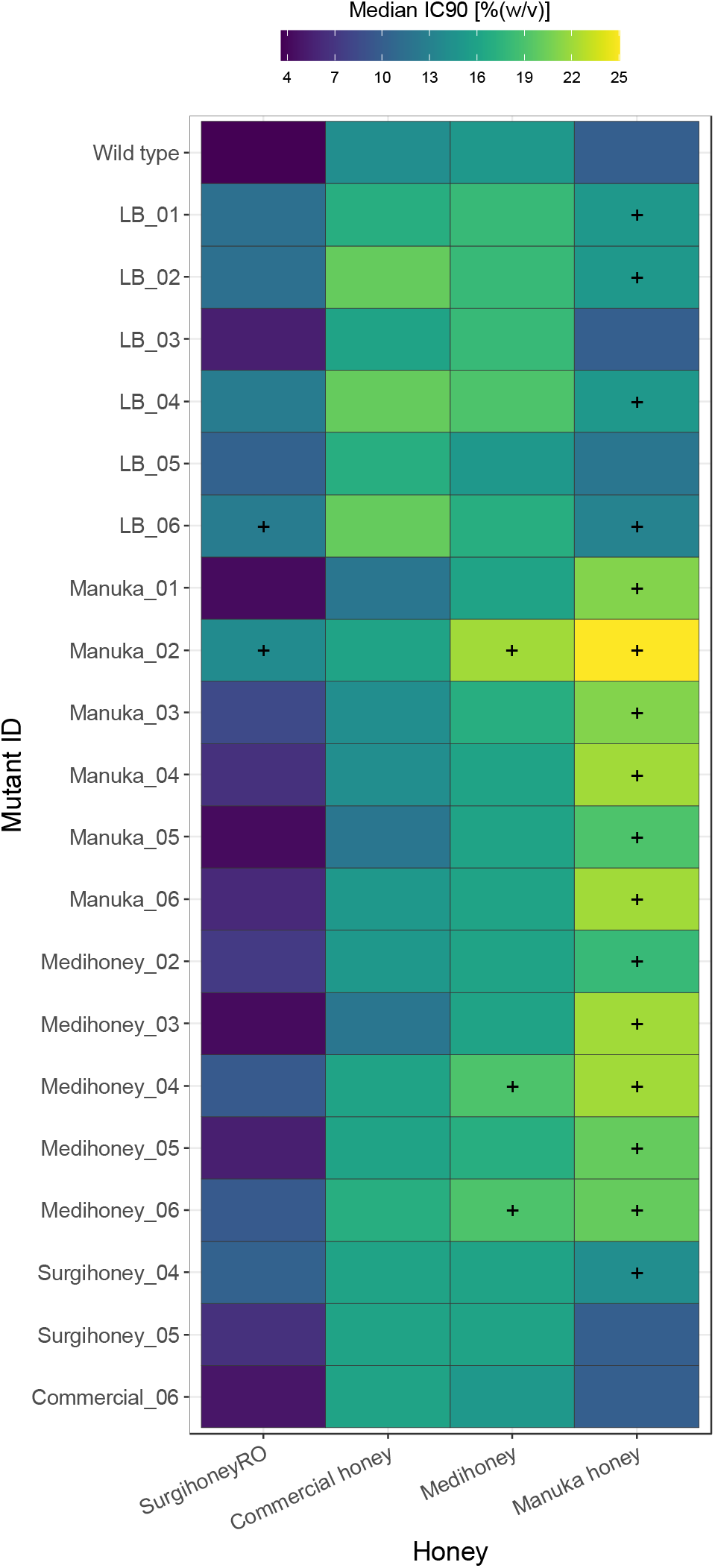
Susceptibility of serially-passaged, putative resistant mutants with four different honey compounds. Each cell gives the median IC_90_ for a given isolate (the ancestral strain K-12 MG1655, in the top row, or a putative resistant mutant serially passaged with one of the four honey compounds, labelled according to compound and replicate selection line) assayed with one of the four honey compounds (columns). Each value is the median of four independent replicates; combinations where all replicates of a putative resistant mutant were higher than all replicates of the ancestor in the same treatment are indicated with a “+”. Individual replicates for each strain are shown in Fig. S4.

### Honey Adaptation Is Linked to Mutations in *nemAR* and *clpP*

Genomic sequencing of our serially passaged isolates revealed changes at several loci (Fig. 2, Table S2). Some loci were mutated multiple times independently in honey-evolved colony isolates, but not in control-evolved isolates (Table S2, Fig. 2), indicating a possible role in adaptation to honey. Two such genes were *clpP* (serine protease, mutated in two Manuka honey-evolved isolates) and *nemR* (DNA-binding transcriptional repressor, mutated in four Manuka honey-evolved isolates and three Medihoney™-evolved isolates; Table S2, Fig. 2). The intergenic region between *ydhL* and *nemR* was mutated in the remaining two Manuka honey-adapted isolates and the remaining two Medihoney™-adapted isolates (Table S2, Fig. 2). An additional gene for which we found mutations in multiple isolates serially passaged in Manuka honey is *yafS* (methyltransferase; Table S2, Fig. 2), located next to *gloB*, the gene encoding glyoxalase II, an enzyme involved in detoxification of reactive aldehydes (Ozyamak *et al*., 2010). In control-evolved isolates, we found parallel mutations in *fimE*, a recombinase responsible for on-to-off switching of type 1 fimbriae expression, and *flhD*, one of two transcriptional activators of the *E. coli* flagellar regulon but also involved in cell division (Klemm, 1986; Iino *et al*., 1988; Liu and Matsumura, 1994; Prüß and Matsumura, 1996), consistent with past work with LB-adapted isolates (Knöppel *et al*., 2018). In summary, we found parallelisms in both control-evolved and honey-evolved isolates, and in particular every Manuka honey- and Medihoney™-evolved isolate had a mutation in or affecting *nemR*. As we discuss below, this may reflect the role of *nemR* in methylglyoxal degradation (Ozyamak *et al*., 2013), an active component of *Leptospermum* honeys.

**Figure 2.**
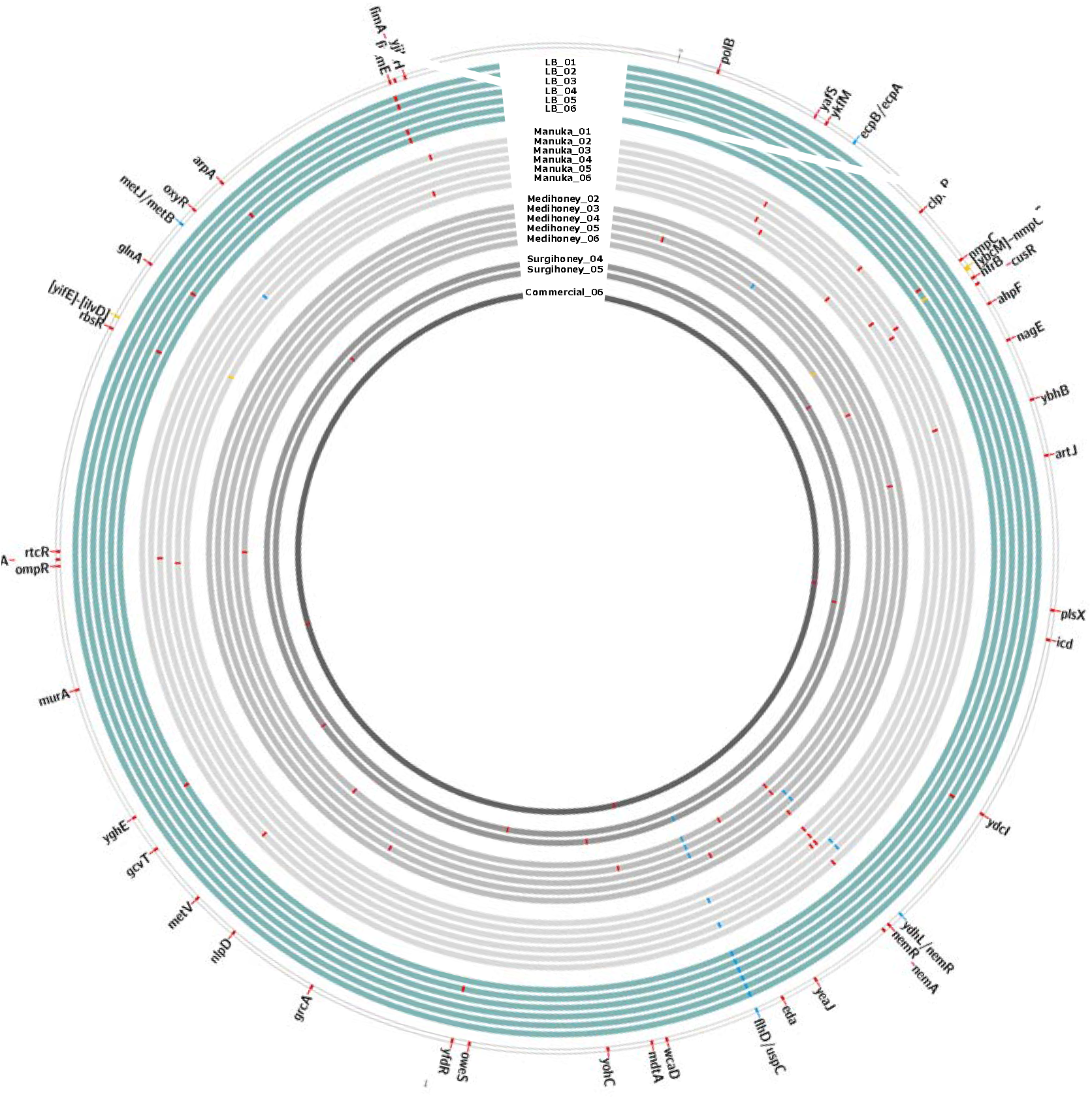
Genetic changes in 20 serially-passaged isolates of *E. coli* K-12 MG1655. The outermost ring represents the ancestral isolate with coloured tiles for all genes in which we found mutations in serially-passaged isolates. Each inner circle represents one serially-passaged isolate: in turquoise: control treatment; in grey: four different honey treatments. Genetic mutations are indicated with coloured tiles: red: mutations in one gene; yellow: intergenetic mutations; blue: deletions >1000bp.

### Single-Gene Knockouts Support a Role for *nemR* in Honey Resistance

We used single-gene knockout variants of several genes that were mutated in serially-passaged isolates to test for further evidence that they play a role in honey resistance (Fig. 3 & S5). The knockout variant *ΔnemR* had increased Manuka honey resistance (all replicates higher than all replicates of the wild-type strain). This variant also showed increased average resistance to Medihoney™, although this was not consistent across all replicate assays (Fig. 3 & S5). This is consistent with the above finding that all isolates serially passaged in Manuka honey or Medihoney™ had mutations in or close to *nemR*, and all were more resistant to Manuka honey and Medihoney™ than the ancestral strain. In Δ*clpP*, the knockout of the other gene for which we observed independent mutations in multiple honey-evolved isolates, we observed a similar pattern: increased resistance to Manuka honey (in all replicate assays) and higher median Medihoney™ resistance (two out of three replicates higher than wild-type). This is consistent with the increased resistance for these honeys we observed with the two Manuka honey-selected isolates with mutations in *clpP* (Fig. 2 & S4). Two other knockout variants, Δ*nemA* and Δ*eda*, had consistently altered resistance to Manuka honey, with Δ*eda* having a higher susceptibility than the ancestral strain. In summary, independent deletion of two genes, *nemR* and *clpP*, which were also directly or indirectly affected by mutations in several of our serially-passaged isolates, conferred increased resistance to Manuka honey and Medihoney™.

**Figure 3.**
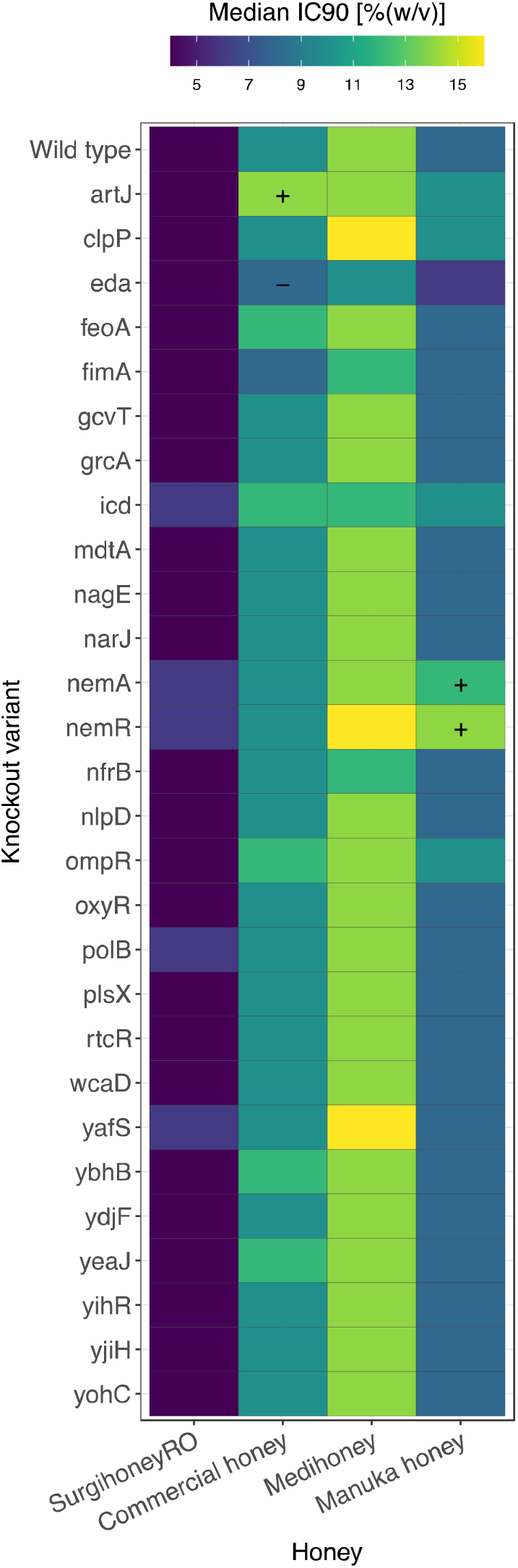
Honey susceptibility of single-gene deletion variants. Each cell shows the median IC_90_ towards four different honeys (columns) for the ancestral strain *E. coli* K-12 BW25113 (top row) or one of 28 single-gene knockout variants (other rows). Each cell is the median value of three independent replicates; combinations where all replicates of a putative resistant mutant were higher/lower than all replicates of the ancestor in the same treatment are indicated with a “+”/“-”. Individual replicates for each strain are shown in Fig. S5.

**Figure 4.**
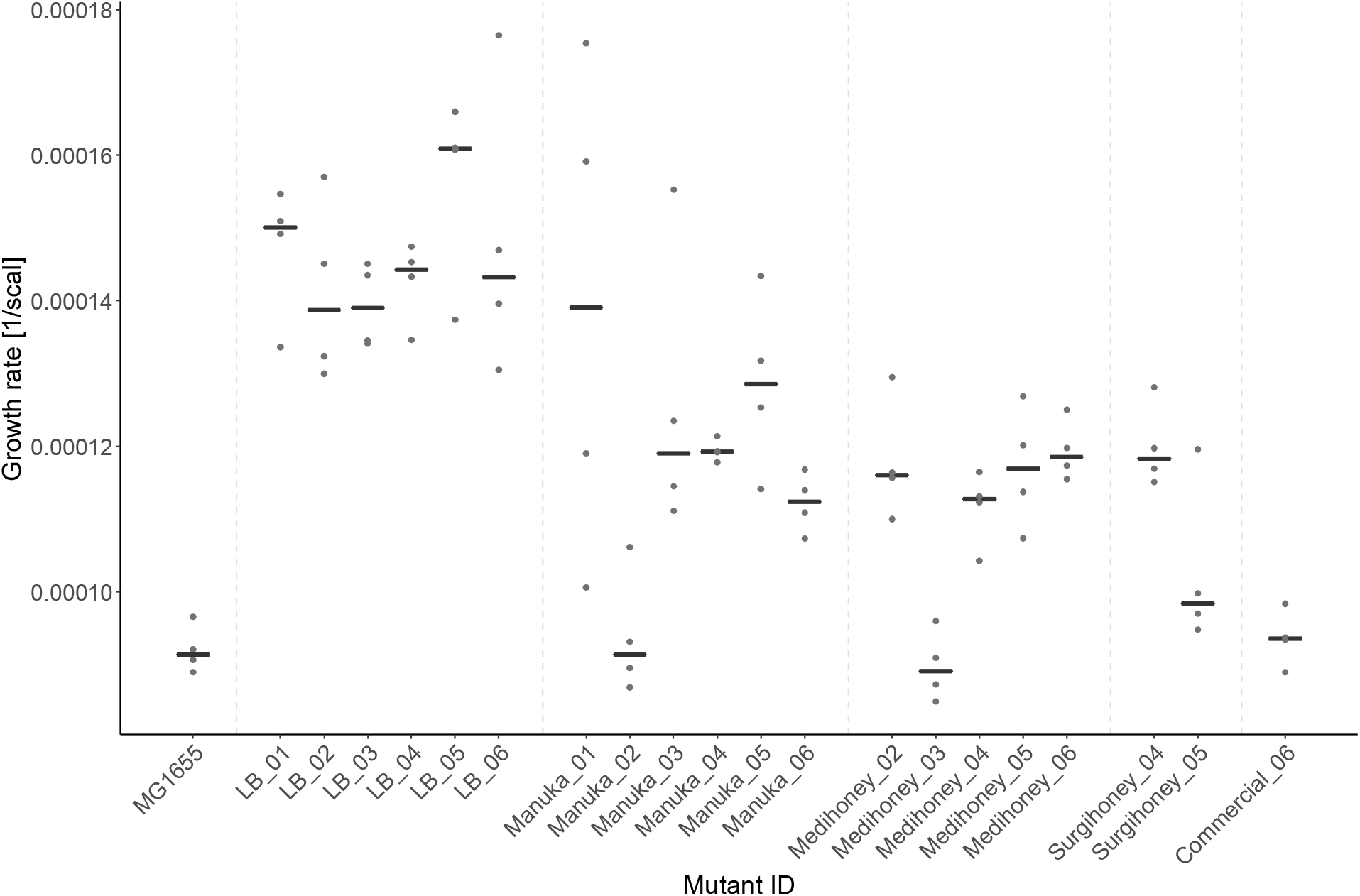
Growth rate of experimentally evolved isolates in the absence of honey. Population growth rate (*y*-axis) is shown for the wild type (MG1655), six mutants serially passaged in honey-free LB (LB_01-LB_06), and 14 mutants serially passaged in different honeys (labelled according to honey and replicate; *x*-axis). Each black line gives the median of four replicates (replicates shown as dots).

### Impaired Growth in the Absence of Honey for Honey-Evolved Compared With Control-Evolved Isolates

Most serially passaged isolates (both honey-evolved and control-evolved) had increased population growth rates relative to the ancestral strain in the absence of honey (Fig. 6). However, when we compared control-evolved isolates with honey-evolved isolates, control-evolved isolates had higher growth rates on average compared to all four types of honey-evolved isolates (linear mixed-effects model: Manuka honey: t_20_ = −4.355, p < 0.001; Medihoney™: t_20_ = −5.538, p < 0.001; SurgihoneyRO™: t_20_ = −4.078, p < 0.001; commercial honey: t_20_ = −4.671, p < 0.001). By contrast, we found little effect of serial passaging on growth yield in evolved isolates compared to the ancestral strain (Fig. S6). Comparing the growth yield in different evolution environments, we found no significant difference between isolates serially passaged in LB and honey-evolved isolates (linear mixed-effects model: all p > 0.05; Fig. S6). In summary, for most experimentally evolved isolates we found a positive effect of serial passage on growth rate, but honey-adapted isolates had a lower growth rate on average compared to isolates serially passaged in the absence of honey.

### No Evidence for Cross-Resistance Between Honey and Antibiotics

We determined the phenotypic resistance of our serially-passaged isolates and the ancestral strain to nine antibiotics of six different classes (Fig. 5; Fig. S7). We found only a single case out of 180 combinations (20 serially-passaged isolates × 9 antibiotics) where antibiotic resistance was consistently altered compared to the ancestral strain (higher/lower IC_90_ for all mutant replicates compared to all replicates of ancestor). This was for one Manuka honey-evolved isolate, Manuka_05, tested with polymyxin B. For seven of the other eight antibiotics, the largest difference shown by individual replicates relative to the median of the ancestral replicates was two-fold or less. For the remaining antibiotic, kanamycin, some individual replicates had differences of four-fold compared to the ancestor. The overall lack of major differences between evolved isolates and the ancestral strain was supported by Analysis of Variance (testing each evolved-isolate-versus-wild-type combination separately, with genotype (evolved vs wild type) and assay compound (antibiotic) as factors, including the interaction term and with *p*-values adjusted using the Holm-Bonferroni method); this indicated a single significant effect of genotype, for isolate Manuka_06. Thus, we found no evidence for appreciable changes in sensitivity to antibiotics in serially-passaged honey-adapted isolates.

**Figure 5.**
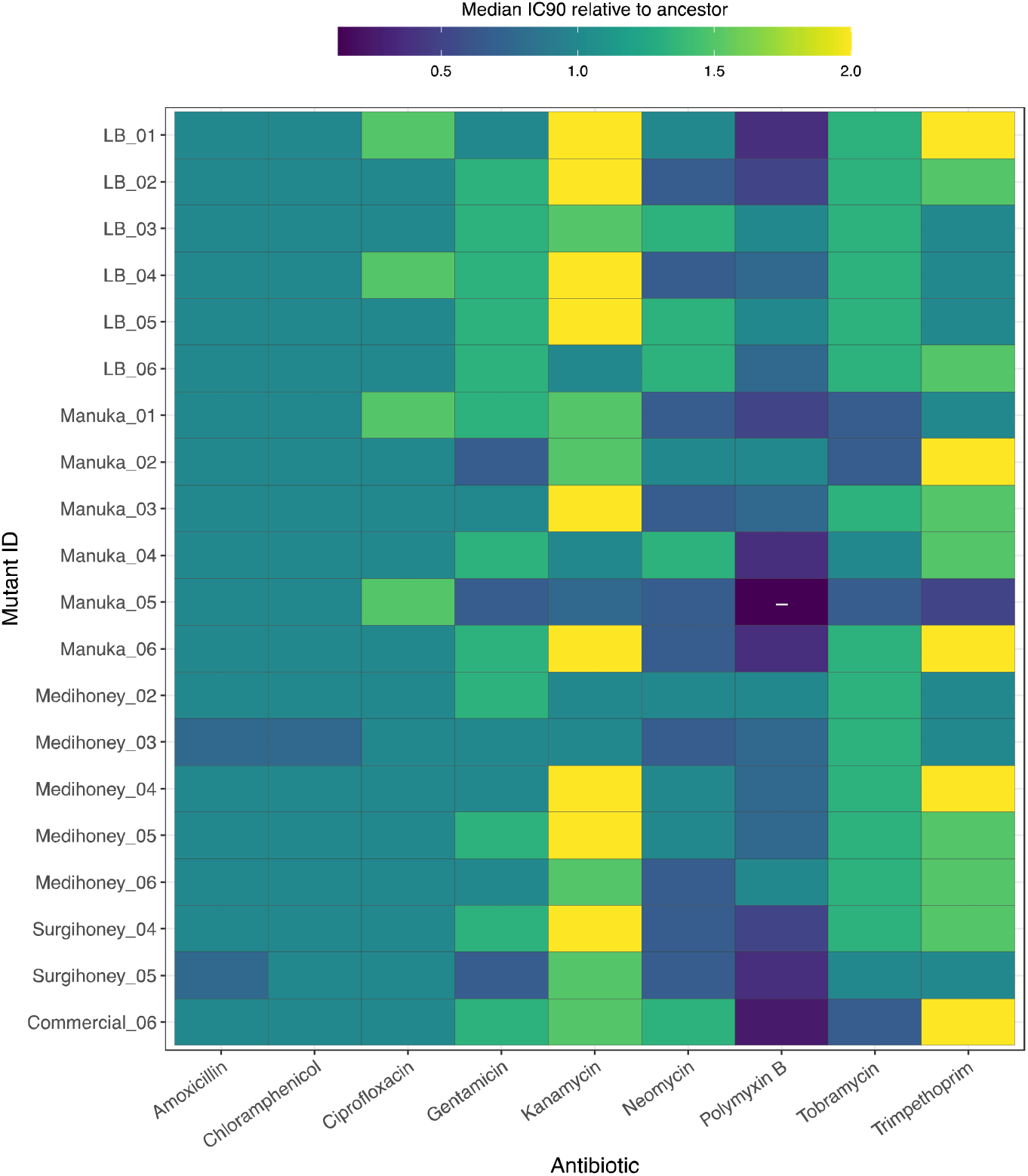
Resistance of serially-passaged, putative honey-resistant mutants against nine different antibiotics, relative to the wild type. The matrix shows the median IC_90_ towards nine antibiotics of 20 serially-passaged isolates relative to the median IC_90_ of the parental strain, *E. coli* K-12 MG1655. Blue/dark seagreen signify susceptibility, yellow/light green resistance relative to the parental strain; combinations where all replicates of a given isolate were lower than all replicates of the ancestor in the same treatment are indicated with a “-”. Individual replicates for each strain are shown in Fig. S7.

### Single-Step Screening Supports Similar Variation of Resistance Evolution Among Honey Products

We tested whether our finding that resistance evolution was more pervasive with Manuka honey than the other honey compounds also holds true when bacteria are exposed to inhibitory honey concentrations suddenly, rather than gradually. We did this by plating aliquots from multiple independent, replicate overnight cultures directly onto honey-supplemented agar, and picking the resulting colony isolates that showed a putative resistance phenotype. As for our serially passaged isolates above, we observed the largest increases in resistance for colony isolates from the Manuka-honey treatment, when assayed with Manuka honey (Fig. S8). For SurgihoneyRO™ and Medihoney™, we observed colonies on some of our plates, but after picking six colonies for each compound and testing their phenotypic resistance, none of them had consistently altered resistance to the honey they were selected with. When we streaked out frozen stocks of these 12 putative honey-resistant colony isolate on honey-supplemented agar at a concentration inhibitory to the parental strain, only 3 out of 12 isolates formed colonies (Medihoney_10, Surgihoney_09, Surgihoney_10). Thus, most putative SurgihoneyRO™- and Medihoney™-resistant mutants showed unstable phenotypic resistance across multiple rounds of isolation, culturing and restreaking (we discuss this further in the context of earlier, similar observations below). For commercial honey, we obtained no viable colonies despite repeating the assay an additional three times on different days (216 aliquots in total for commercial honey). In summary, we isolated single-step resistant mutants with Manuka honey, but found no colony isolates with robust resistance phenotypes to SurgihoneyRO™, Medihoney™ or commercial honey, consistent with our finding in serial passage that resistance evolved most readily against Manuka honey.

## DISCUSSION

We found *E. coli* does not easily develop large increases in honey resistance upon serial passage at gradually increasing concentrations, or upon plating of many replicate populations at inhibitory concentrations. However, we identified a set of genes important in adaptation to honey, which conferred moderate increases in resistance to Manuka honey. The known physiological role of this genetic pathway helps to explain why these genetic changes, and associated increases in phenotypic resistance, were specific to *Leptospermum* honeys (detoxification of methylglyoxal, which has been found to be the major contributor to the antibacterial activity of these honeys (Adams *et al*., 2008; Mavric *et al*., 2008)). This indicates the likelihood of honey resistance upon medical application may depend critically on the type or combination of honey products used. Furthermore, we found that honey adaptation *in vitro* had only minimal side-effects for antibiotic resistance.

The first important implication of our results is that *E. coli* does not readily become resistant to honey, which is promising in the context of expanding medical use of honey. Previous studies also had difficulty in isolating mutants with stable honey resistance, and/or found that honey resistance phenotypes revert quickly in the absence of honey (Cooper *et al*., 2010). This is consistent with our observation that several putative resistant mutants from our single-step screen had unstable phenotypes, and some serially passaged populations that attained the ability to grow at increased honey concentrations subsequently lost this ability upon transfer to fresh medium (referred to by (Abdel-Azim *et al*., 2019) as *second transfer crash* (STC)). One possible explanation is that non-genetic changes such as persister formation or other morphological changes enable temporary population survival or growth in otherwise inhibitory concentrations. Consistent with honey inducing physiological responses that are not necessarily heritable, Brudzynski and Sjaarda (2014) observed morphological changes in *E. coli* cultures exposed to honey, including changes in cell shape / filamentation.

The second key implication of our results is that, to our knowledge, this is the first report of a specific genetic mechanism linked to decreased honey susceptibility, namely changes affecting the *nemAR* operon, and alternatively via changes affecting *clpP*. In support, *nemR* and its operon have previously been described as being involved in physiological processes that we can expect to be beneficial during honey exposure. Manuka honey and Medihoney™ are both *Leptospermum* honeys, whose high non-peroxide antimicrobial activity is attributed to their relatively high content of methylglyoxal (Allen *et al*., 1991; Mavric *et al*., 2008), a small amount of which bacteria produce intracellularly (Tötemeyer *et al*., 1998). Ozyamak *et al*. (2013) linked the *nemAR* operon to methylglyoxal detoxification. The *nemAR* operon is located upstream from the glyoxalase system (Ozyamak *et al*., 2013) which consists of GlxI and GlxII (glyoxalase I and II, encoded by *gloA* and *gloB* respectively), the two most important enzymes in the detoxification pathway of methylglyoxal (Clugston *et al*., 1998; MacLean *et al*., 1998; Mannervik, 2008). Deletion of *nemR*, the repressor gene, results in increased transcription both of *nemA* and *gloA* (Ozyamak *et al*., 2013). Thus, the finding that all isolates serially passaged in supposedly methylglyoxal-rich *Leptospermum* honeys have mutations in or close to the repressor gene *nemR* allows for the speculation that, in this context, deletion of this gene is beneficial because it leads to increased methylglyoal detoxification. Similarly, *clpP*, another gene we found independently mutated in multiple Manuka honey-adapted isolates, was shown by Jenkins *et al*. (2014) to be overexpressed in *Leptospermum* honey-exposed methicillin-resistant *S. aureus*, supporting a role for this gene in honey resistance in multiple species.

The importance of methylglyoxal detoxification in increased honey resistance is further supported by the variation we observed among honey compounds. Robust resistant mutants emerged in our single-step screen only with a *Leptospermum* honey (Manuka) with a high concentration of methylglyoxal, but not with the other three compounds. During serial passage we saw a similar trend: the most consistent increases in resistance were with the two *Leptospermum* honey products. By contrast, resistance of putative single-step resistant mutants isolated from SurgihoneyRO™-supplemented agar was not stable, and we could not isolate any spontaneous mutants for the commercial honey we used (despite observing increased population growth at inhibitory concentrations during serial passage). This commercial honey is a blend of honeys with unknown flower sources from different South-American countries. Drawing a parallel to antibiotics, where combinations of antibiotics can make it harder for bacteria to evolve resistance (Baym *et al*., 2016), we speculate that resistance evolution against this honey was rare in our experiments because it is a form of combination treatment. This raises the possibility that effective application of honey in treatment may benefit from combining multiple types of honey products with different mechanisms of action. We note that despite variation among honey products, all isolates in our experiment were inhibited by honey concentrations comparable to those found in medical-grade honey products (63–100% in products recommended/licensed for medical wound care, http://www.medihoney.de/index.html). We hypothesize that the multi-faceted nature of honey’s antibacterial activity (Wang *et al*., 2012; Nolan *et al*., 2019) contributes to the difficulty of bacteria evolving resistance to such high concentrations.

The third major implication of our results is that honey adaptation did not come with collateral effects on antibiotic resistance. The lack of collateral effects on antibiotic resistance is an important and promising aspect in the context of wider medical application of honey, and contrasts starkly with the frequent occurrence of cross-resistances between antibiotics (Szybalski and Bryson, 1952; Gutmann *et al*., 1985) and between antibiotics and some other antimicrobial agents (Loughlin *et al*., 2002; Braoudaki and Hilton, 2004; Baker-Austin *et al*., 2006; Allen *et al*., 2017; Bischofberger *et al*., 2020). We do not exclude that more significant changes in antibiotic resistance might be observed in mutants or strains with larger changes in honey resistance than the moderate increases we observed here, if such strains exist. A key avenue for future work is therefore to determine whether any genetic variation in natural populations associated with variable honey resistance is independent of variation in antibiotic resistance, as in our experiment. Another important aspect to consider is honey’s potential effect on other bacterial virulence factors. Biofilm formation is an effective bacterial defense mechanism against a wide range of antimicrobials (Costerton *et al*., 1995). Several of the genes we found mutated in honey-adapted isolates have a known role in biofilm formation (*fimA, fimE, nlpD, ompR, yeaJ* (Prigent-Combaret *et al*., 2001; Niba *et al*., 2007; Wu and Outten, 2009; Amores *et al*., 2017)). Honey or honey-resistance mechanisms might therefore also have downstream effects for biofilm formation or the expression of other virulence factors. On the other hand, because honey has shown effective inhibition of a wide range of pathogens (Carter *et al*., 2016; Hillitt *et al*., 2017; Yabes *et al*., 2017), and the principal genes involved in honey resistance in our experiment are conserved in multiple pathogenic species (e.g., *nemR* or analogues are present in *Klebsiella pneumonia, Acinetobacter baumannii* and *Salmonella typhimurium*), our work identifies candidate loci that may be involved in resistance in other species. This is another relevant area for future research in the context of medical honey application.

Our results also provided indirect evidence of growth costs associated with honey adaptation. Growth in the absence of selecting antibacterials is widely considered an important parameter in the long-term spread of resistance (Andersson and Hughes, 2010). The larger increase in growth rate observed in our control-evolved isolates compared to honey-adapted isolates suggests either (i) that honey adaptation constrained adaptation to other aspects of the environment, preventing acquisition of other beneficial mutations, or (ii) that honey-resistance mutations combined with other types of beneficial mutations conferred a smaller net increase in growth rate in honey-free growth medium compared to other beneficial mutations alone. Our sequence data are more consistent with the former: control-evolved populations had parallel mutations in flagella and fimbriae genes (*flhD* and *fimE*), consistent with past work with LB-adapted *E. coli* (Knöppel *et al*., 2018). These mutations were much less common in honey-adapted colony isolates (Fig. 2). This could be due, for example, to epistatic interactions between the different types of adaptive mutations (Scanlan *et al*., 2015). The parallelism we observed for these genes might also be responsible for the increased honey resistance of control isolates (Fig. S4). However, a full understanding of the growth costs and benefits associated with individual honey-resistance alleles and their interactions with other types of beneficial mutations is beyond the scope of this paper, and was not our aim here, but would make an interesting avenue for future work.

In conclusion, honey resistance in our experiment only evolved with a subset of the compounds we tested, and only to a moderate degree. This is promising in the context of medical application of honey. A further positive aspect is that we found no evidence of strong downstream effects on antibiotic susceptibility in isolates adapted to honey *in vitro* via chromosomal mutation. Finally, we identified putative genetic mechanisms involved in honey adaptation, via changes affecting genes involved in detoxifying methylglyoxal, making this mechanism most relevant for *Leptospermum* honeys such as Manuka honey.

## Acknowledgments

A.B. thanks Richard Allen for help with statistical analysis. A.H. acknowledges Swiss National Science Foundation project 31003A_165803.

## Data Archiving Statement

The data that support the findings of this study will be made openly available at Dryad Digital Repository: to be completed after manuscript is accepted for publication.

## References

Abdel-Azim, S.G., Abdel-Azim, A.G., Piasecki, B.P., and Abdel-Azim, G.A. (2019) Characterization of the Gain and Loss of Resistance to Antibiotics versus Tolerance to Honey as an Antimutagenic and Antimicrobial Medium in Extended-Time Serial Transfer Experiments. Pharmacognosy Res 11: 147–54.

Adams, C.J., Boult, C.H., Deadman, B.J., Farr, J.M., Grainger, M.N.C., Manley-Harris, M., and Snow, M.J. (2008) Isolation by HPLC and characterisation of the bioactive fraction of New Zealand manuka (Leptospermum scoparium) honey. Carbohydr Res 343: 651–659.

Allen, K.L., Molan, P.C., and Reid, G.M. (1991) A Survey of the Antibacterial Activity of Some New Zealand Honeys. J Pharm Pharmacol 43: 817–822.

Allen, R.C., Pfrunder-Cardozo, K.R., Meinel, D., Egli, A., and Hall, A.R. (2017) Associations among antibiotic and phage resistance phenotypes in natural and clinical Escherichia coli isolates. MBio 8: e01341–17.

Amores, G.R., De Las Heras, A., Sanches-Medeiros, A., Elfick, A., and Silva-Rocha, R. (2017) Systematic identification of novel regulatory interactions controlling biofilm formation in the bacterium Escherichia coli. Sci Rep 7: 1–14.

Andersson, D.I. and Hughes, D. (2010) Antibiotic resistance and its cost: Is it possible to reverse resistance? Nat Rev Microbiol 8: 260–271.

Ankley, L.M., Monteiro, M.P., Camp, K.M., O’Quinn, R., and Castillo, A.R. (2020) Manuka honey chelates iron and impacts iron regulation in key bacterial pathogens. J Appl Microbiol 128: 1015–1024.

Baba, T., Ara, T., Hasegawa, M., Takai, Y., Okumura, Y., Baba, M., et al. (2006) Construction of Escherichia coli K-12 in-frame, single-gene knockout mutants: The Keio collection. Mol Syst Biol 2:.

Badet, C. and Quero, F. (2011) The in vitro effect of manuka honeys on growth and adherence of oral bacteria. Anaerobe 17: 19–22.

Baker-Austin, C., Wright, M.S., Stepanauskas, R., and McArthur, J. V. (2006) Co-selection of antibiotic and metal resistance. Trends Microbiol 14: 176–182.

Basualdo, C., Sgroy, V., Finola, M.S., and Marioli, J.M. (2007) Comparison of the antibacterial activity of honey from different provenance against bacteria usually isolated from skin wounds. Vet Microbiol 124: 375–381.

Baym, M., Stone, L.K., and Kishony, R. (2016) Multidrug evolutionary strategies to reverse antibiotic resistance. Science (80-) 351:.

Bischofberger, A.M., Baumgartner, M., Pfrunder-Cardozo, K.R., Allen, R.C., and Hall, A.R. (2020) Associations between sensitivity to antibiotics, disinfectants and heavy metals in natural, clinical and laboratory isolates of Escherichia coli. Environ Microbiol 00: 1–16.

Bolger, A.M., Lohse, M., and Usadel, B. (2014) Trimmomatic: A flexible trimmer for Illumina sequence data. Bioinformatics 30: 2114–2120.

Braoudaki, M. and Hilton, A.C. (2004) Low level of cross-resistance between triclosan and antibiotics in Escherichia coli K-12 and E. coli O55 compared to E. coli O157. FEMS Microbiol Lett 235: 305–309.

Breasted, J.H. (1948) The Edwin Smith Surgical Papyrus, Breasted, J.H. (ed) Chicago: The University of Chicago Press.

Brudzynski, K. and Sjaarda, C. (2014) Antibacterial compounds of canadian honeys target bacterial cell wall inducing phenotype changes, growth inhibition and cell lysis that resemble action of ß-lactam antibiotics. PLoS One 9: 106967.

Camplin, A.L. and Maddocks, S.E. (2014) Manuka honey treatment of biofilms of Pseudomonas aeruginosa results in the emergence of isolates with increased honey resistance. Ann Clin Microbiol Antimicrob 13: 19.

Carnwath, R., Graham, E.M., Reynolds, K., and Pollock, P.J. (2014) The antimicrobial activity of honey against common equine wound bacterial isolates. Vet J 199: 110–114.

Carter, D.A., Blair, S.E., Cokcetin, N.N., Bouzo, D., Brooks, P., Schothauer, R., and Harry, E.J. (2016) Therapeutic manuka honey: No longer so alternative. Front Microbiol 7: 1–11.

Chittezham Thomas, V., Kinkead, L.C., Janssen, A., Schaeffer, C.R., Woods, K.M., Lindgren, J.K., et al. (2013) A dysfunctional tricarboxylic acid cycle enhances fitness of Staphylococcus epidermidis during ß-lactam stress. MBio 4: e00437–13.

Clugston, S.L., Barnard, J.F.J., Kinach, R., Miedema, D., Ruman, R., Daub, E., and Honek, J.F. (1998) Overproduction and characterization of a dimeric non-zinc glyoxalase I from Escherichia coli: Evidence for optimal activation by nickel ions. Biochemistry 37: 8754–8763.

Cooke, J., Dryden, M., Patton, T., Brennan, J., and Barrett, J. (2015) The antimicrobial activity of prototype modified honeys that generate reactive oxygen species (ROS) hydrogen peroxide. BMC Res Notes 8: 20.

Cooper, R.A., Jenkins, L., Henriques, A.F.M., Duggan, R.S., and Burton, N.F. (2010) Absence of bacterial resistance to medical-grade manuka honey. Eur J Clin Microbiol Infect Dis 29: 1237–1241.

Costerton, J.W., Lewandowski, Z., Caldwell, D.E., Korber, D.R., and Lappin-Scott, H.M. (1995) Microbial biofilms. Annu Rev Microbiol 49: 711–45.

Deatherage, D.E. and Barrick, J.E. (2014) Identification of mutations in laboratory-evolved microbes from next-generation sequencing data using breseq. Methods Mol Biol 1151: 165–188.

Descottes, B. (2009) Cicatrisation par le miel, l’expérience de 25 années. Phytotherapie 7: 112–116.

Dowd, S.E., Sun, Y., Secor, P.R., Rhoads, D.D., Wolcott, B.M., James, G.A., and Wolcott, R.D. (2008) Survey of bacterial diversity in chronic wounds using Pyrosequencing, DGGE, and full ribosome shotgun sequencing. BMC Microbiol 8: 43.

Dunford, C.E. and Hanano, R. (2004) Acceptability to patients of a honey dressing for non-healing venous leg ulcers. J Wound Care 13: 193–197.

Gutmann, L., Williamson, R., Moreau, N., Kitzis, M.D., Collatz, E., Acar, J.F., and Goldstein, F.W. (1985) Cross-resistance to nalidixic acid, trimethoprim, and chloramphenicol associated with alterations in outer membrane proteins of klebsiella, enterobacter, and serratia. J Infect Dis 151: 501–507.

Haffejee, I.E. and Moosa, A. (1985) Honey in the treatment of infantile gastroenteritis. Br Med J (Clin Res Ed) 290: 1866.

Halstead, F.D., Webber, M.A., Rauf, M., Burt, R., Dryden, M., and Oppenheim, B.A. (2016) In vitro activity of an engineered honey, medical-grade honeys, and antimicrobial wound dressings against biofilm-producing clinical bacterial isolates. J Wound Care 25: 93–102.

Henriques, A.F., Jenkins, R.E., Burton, N.F., and Cooper, R.A. (2011) The effect of manuka honey on the structure of Pseudomonas aeruginosa. Eur J Clin Microbiol Infect Dis 30: 167–171.

Hillitt, K.L., Jenkins, R.E., Spiller, O.B., and Beeton, M.L. (2017) Antimicrobial activity of Manuka honey against antibiotic-resistant strains of the cell wall-free bacteria Ureaplasma parvum and Ureaplasma urealyticum. Lett Appl Microbiol 64: 198–202.

Iino, T., Komeda, Y., Kutsukake, K., Macnab, R.M., Matsumura, P., Parkinson, J.S., et al. (1988) New unified nomenclature for the flagellar genes of Escherichia coli and Salmonella typhimurium. Microbiol Rev 52: 533–535.

Irish, J., Blair, S., and Carter, D.A. (2011) The antibacterial activity of honey derived from Australian flora. PLoS One 6: e18229.

Jenkins, R., Burton, N., and Cooper, R. (2014) Proteomic and genomic analysis of methicillin-resistant staphylococcus aureus (MRSA) exposed to manuka honey in vitro demonstrated down-regulation of virulence markers. J Antimicrob Chemother 69: 603–615.

Klemm, P. (1986) Two regulatory fim genes, fimB and fimE, control the phase variation of type 1 fimbriae in Escherichia coli. EMBO J 5: 1389–1393.

Knipping, S., Grünewald, B., and Hirt, R. (2012) Erste Erfahrungen mit medizinischem Honig in der Wundbehandlung im Kopf-Hals-Bereich. HNO 60: 830–836.

Knöppel, A., Knopp, M., Albrecht, L.M., Lundin, E., Lustig, U., Näsvall, J., and Andersson, D.I. (2018) Genetic adaptation to growth under laboratory conditions in Escherichia coli and Salmonella enterica. Front Microbiol 9: 1–16.

Knottenbelt, D.C. (2014) Honey in wound management: Myth, mystery, magic or marvel? Vet J 199: 5–6.

Kronda, J.M., Cooper, R.A., and Maddocks, S.E. (2013) Manuka honey inhibits siderophore production in Pseudomonas aeruginosa. J Appl Microbiol 115: 86–90.

Kwakman, P.H.S., Van den Akker, J.P.C., Güçlü, A., Aslami, H., Binnekade, J.M., de Boer, L., et al. (2008) Medical Grade Honey Kills Antibiotic□Resistant Bacteria In Vitro and Eradicates Skin Colonization. Clin Infect Dis 46: 1677–1682.

Kwakman, P.H.S., te Velde, A.A., de Boer, L., Vandenbroucke-Grauls, C.M.J.E., and Zaat, S.A.J. (2011) Two major medicinal honeys have different mechanisms of bactericidal activity. PLoS One 6: e17709.

Lee, J.H., Park, J.H., Kim, J.A., Neupane, G.P., Cho, M.H., Lee, C.S., and Lee, J. (2011) Low concentrations of honey reduce biofilm formation, quorum sensing, and virulence in Escherichia coli O157:H7. Biofouling 27: 1095–1104.

Liu, X. and Matsumura, P. (1994) The FlhD/FlhC complex, a transcriptional activator of the Escherichia coli flagellar class II operons. J Bacteriol 176: 7345–7351.

Loughlin, M.F., Jones, M. V., and Lambert, P.A. (2002) Pseudomonas aeruginosa cells adapted to benzalkonium chloride show resistance to other membrane-active agents but not to clinically relevant antibiotics. J Antimicrob Chemother 49: 631–639.

Lu, J., Carter, D.A., Turnbull, L., Rosendale, D., Hedderley, D., Stephens, J., et al. (2013) The Effect of New Zealand Kanuka, Manuka and Clover Honeys on Bacterial Growth Dynamics and Cellular Morphology Varies According to the Species. PLoS One 8: 55898.

Lu, J., Cokcetin, N.N., Burke, C.M., Turnbull, L., Liu, M., Carter, D.A., et al. (2019) Honey can inhibit and eliminate biofilms produced by Pseudomonas aeruginosa. Sci Rep 9: 1–13.

MacLean, M.J., Ness, L.S., Ferguson, G.P., and Booth, I.R. (1998) The role of glyoxalase I in the detoxification of methylglyoxal and in the activation of the KefB K+ efflux system in Escherichia coli. Mol Microbiol 27: 563–571.

Maddocks, S.E. and Jenkins, R.E. (2013) Honey: A sweet solution to the growing problem of antimicrobial resistance? Future Microbiol 8: 1419–1429.

Magnet, S. and Blanchard, J.S. (2005) Molecular insights into aminoglycoside action and resistance. Chem Rev 105: 477–497.

Mannervik, B. (2008) Molecular enzymology of the glyoxalase system. Drug Metabol Drug Interact 23: 13–27.

Mavric, E., Wittmann, S., Barth, G., and Henle, T. (2008) Identification and quantification of methylglyoxal as the dominant antibacterial constituent of Manuka (Leptospermum scoparium) honeys from New Zealand. Mol Nutr Food Res 52: 483–489.

Merckoll, P., Jonassen, T.Ø., Vad, M.E., Jeansson, S.L., and Melby, K.K. (2009) Bacteria, biofilm and honey: A study of the effects of honey on “planktonic” and biofilm-embedded chronic wound bacteria. Scand J Infect Dis 41: 341–347.

Molan, P.C. (1992) THE ANTIBACTERIAL ACTIVITY OF HONEY. 1. The nature of the antibacterial activity. Bee World1 73: 5–28.

Molan, P.C. (1999) Why honey is effective as a medicine. I. Its use in modern medicine. Bee World 80: 80–92.

Molan, P.C. and Betts, J.A. (2004) Clinical usage of honey as a wound dressing: an update. J Wound Care 13: 353–356.

Niba, E.T.E., Naka, Y., Nagase, M., Mori, H., and Kitakawa, M. (2007) A genome-wide approach to identify the genes involved in biofilm formation in E. coli. DNA Res 14: 237–246.

Nolan, V.C., Harrison, J., and Cox, J.A.G. (2019) Dissecting the antimicrobial composition of honey. Antibiotics 8: 1–16.

O’Neill, J. (2014) Antimicrobial Resistance: Tackling a crisis for the health and wealth of nations The Review on Antimicrobial Resistance Chaired.

Oliveira, A., Ribeiro, H.G., Silva, A.C., Silva, M.D., Sousa, J.C., Rodrigues, C.F., et al. (2017) Synergistic antimicrobial interaction between honey and phage against Escherichia coli biofilms. Front Microbiol 8: 2407.

Oliveira, A., Sousa, J.C., Silva, A.C., Melo, L.D.R., and Sillankorva, S. (2018) Chestnut Honey and Bacteriophage Application to Control Pseudomonas aeruginosa and Escherichia coli Biofilms: Evaluation in an ex vivo Wound Model. Front Microbiol 9: 1725.

Ozyamak, E., De Almeida, C., De Moura, A.P.S., Miller, S., and Booth, I.R. (2013) Integrated stress response of Escherichia coli to methylglyoxal: Transcriptional readthrough from the nemRA operon enhances protection through increased expression of glyoxalase I. Mol Microbiol 88: 936–950.

Ozyamak, E., Black, S.S., Walker, C.A., MacLean, M.J., Bartlett, W., Miller, S., and Booth, I.R. (2010) The critical role of S-lactoylglutathione formation during methylglyoxal detoxification in Escherichia coli. Mol Microbiol 78: 1577–1590.

Paulsen, I.T., Brown, M.H., and Skurray, R.A. (1996) Proton-dependent multidrug efflux systems. Microbiol Rev 60: 575–608.

Philippon, A., Labia, R., and Jacoby, G. (1989) MINIREVIEW Extended-Spectrum beta-Lactamases. Antimicrob Agents Chemother 33: 1131–1136.

Prigent-Combaret, C., Brombacher, E., Vidal, O., Ambert, A., Lejeune, P., Landini, P., and Dorel, C. (2001) Complex regulatory network controls initial adhesion and biofilm formation in Escherichia coli via regulation of the csgD gene. J Bacteriol 183: 7213–7223.

Prüß, B.M. and Matsumura, P. (1996) A regulator of the flagellar regulon of Escherichia coli, flhD, also affects cell division. J Bacteriol 178: 668–674.

Scanlan, P.D., Hall, A.R., Blackshields, G., Friman, V.P., Davis, M.R., Goldberg, J.B., and Buckling, A. (2015) Coevolution with bacteriophages drives genome-wide host evolution and constrains the acquisition of abiotic-beneficial mutations. Mol Biol Evol 32: 1425–1435.

Shan, Y., Lazinski, D., Rowe, S., Camilli, A., and Lewis, K. (2015) Genetic Basis of Persister Tolerance to Aminoglycosides in Escherichia coli. MBio 6: e00078–15.

Su, Y. bin, Peng, B., Li, H., Cheng, Z. xue, Zhang, T. tuo, Zhu, J. xin, et al. (2018) Pyruvate cycle increases aminoglycoside efficacy and provides respiratory energy in bacteria. Proc Natl Acad Sci U S A 115: E1578–E1587.

Suojala, L., Kaartinen, L., and Pyörälä, S. (2013) Treatment for bovine Escherichia coli mastitis - an evidence-based approach. J Vet Pharmacol Ther 36: 521–531.

Szybalski, W. and Bryson, V. (1952) Genetic studies on microbial cross resistance to toxic agents: Cross resistance of Escherichia coli to fifteen antibiotics. J Bacteriol 64: 489–499.

Tadesse, D.A., Zhao, S., Tong, E., Ayers, S., Singh, A., Bartholomew, M.J., and McDermott, P.F. (2012) Antimicrobial drug resistance in Escherichia coli from humans and food animals, United States, 1950-2002. Emerg Infect Dis 18: 741–749.

Tötemeyer, S., Booth, N.A., Nichols, W.W., Dunbar, B., and Booth, I.R. (1998) From famine to feast: The role of methylglyoxal production in Escherichia coli. Mol Microbiol 27: 553–562.

Tramuta, C., Nebbia, P., Robino, P., Giusto, G., Gandini, M., Chiadò-Cutin, S., and Grego, E. (2017) Antibacterial activities of Manuka and Honeydew honey-based membranes against bacteria that cause wound infections in animals. Schweiz Arch Tierheilkd 159: 117–121.

Truchado, P., López-Gálvez, F., Gil, M.I., Tomás-Barberán, F.A., and Allende, A. (2009) Quorum sensing inhibitory and antimicrobial activities of honeys and the relationship with individual phenolics. Food Chem 115: 1337–1344.

Vandamme, L., Heyneman, A., Hoeksema, H., Verbelen, J., and Monstrey, S. (2013) Honey in modern wound care: A systematic review. Burns 39: 1514–1525.

Wang, R., Starkey, M., Hazan, R., and Rahme, L.G. (2012) Honey’s ability to counter bacterial infections arises from both bactericidal compounds and QS inhibition. Front Microbiol 3: 1–8.

Wasfi, R., Elkhatib, W.F., and Khairalla, A.S. (2016) Effects of selected egyptian honeys on the cellular ultrastructure and the gene expression profile of Escherichia coli. PLoS One 11: e0150984.

Werner, A. and Laccourreye, O. (2011) Honey in otorhinolaryngology: When, why and how? Eur Ann Otorhinolaryngol Head Neck Dis 128: 133–137.

White, J.W., Subers, M.H., and Schepartz, A.I. (1963) The identification of inhibine, the antibacterial factor in honey, as hydrogen peroxide and its origin in a honey glucose-oxidase system. BBA - Biochim Biophys Acta 73: 57–70.

WHO (2018) Fact Sheet Antibacterial Resistance.

Willix, D.J., Molan, P.C., and Harfoot, C.G. (1992) A comparison of the sensitivity of wound□infecting species of bacteria to the antibacterial activity of manuka honey and other honey. J Appl Bacteriol 73: 388–394.

Wu, Y. and Outten, F.W. (2009) IscR controls iron-dependent biofilm formation in Escherichia coli by regulating type I fimbria expression. J Bacteriol 191: 1248–1257.

Yabes, J.M., White, B.K., Murray, C.K., Sanchez, C.J., Mende, K., Beckius, M.L., et al. (2017) In Vitro activity of Manuka Honey and polyhexamethylene biguanide on filamentous fungi and toxicity to human cell lines. Med Mycol 55: 334–343.

Zhou, S., Zhuang, Y., Zhu, X., Yao, F., Li, Haiyan, Li, Huifang, et al. (2019) YhjX regulates the growth of Escherichia coli in the presence of a subinhibitory concentration of gentamicin and mediates the adaptive resistance to gentamicin. Front Microbiol 10: 1–10.

