## Supplementary Data for "Evolution of Honey Resistance in Experimental Populations of Bacteria Depends on the Type of Honey, and Has no Major Side Effects for Antibiotic Susceptibility"

**Tables**

**Supplementary Table 1. Range of honey concentrations tested in assays for phenotypic honey resistance.** Range and increment are given in %(w/v).

|  | SurgihoneyRO™ | | Medihoney™ | | Manuka honey | | Commercial honey | |
| --- | --- | --- | --- | --- | --- | --- | --- | --- |
|  | range | increment | range | increment | range | increment | range | increment |
| assay A | 0.8-8 | 0.8 | 1.6-16 | 1.6 | 1.6-16 | 1.6 | 1.6-16 | 1.6 |
| assay B | 1.6-16 | 1.6 | 2.4-24 | 2.4 | 2.4-24 | 2.4 | 2.4-24 | 2.4 |
| assay C | 1.2-13.2 | 1.2 | 4-24 | 2 | 4-24 | 2 | 4-24 | 2 |
| assay D | 2-18 | 2 | 2-18 | 2 | 2-18 | 2 | 2-18 | 2 |

**Supplementary Table 2. Genetic changes identified in isolates serial passaged with/without different honey products.** The table contains all mutations registered in serially-passaged isolates (14 isolates adapted to four different honeys, six control isolates) using the Illumina Hiseq 4000 platform. Gene description according to breseq and the reference sequence of the ancestral isolate, E. coli K-12 MG1655. (*see separate .xslx document “TableS2_HoneyResistance.xslx”*)

**Supplementary Figures**

**Supplementary Figure 1.**

see file *FigureS1_HoneyResistance*

**Supplementary Figure 1. Schematic protocol of experimental resistance evolution.** (A) Serial passage: For each replicate selection line, independent overnight cultures were used to inoculate a range of honey concentrations and honey-free medium (LB). The highest honey concentration that showed viable growth ((OD_600_ 24h – OD_600_ 0h) > 0.1, transfer culture (outlined in red)) was used to inoculate wells filled with various concentrations in a new microplate. Control treatment was performed by daily passaging in honey-free medium (LB). This was repeated daily for 22 days. (B) Single-step screening: for each compound we made multiple overnight cultures before plating on honey-supplemented agar and screening for resistant mutants.

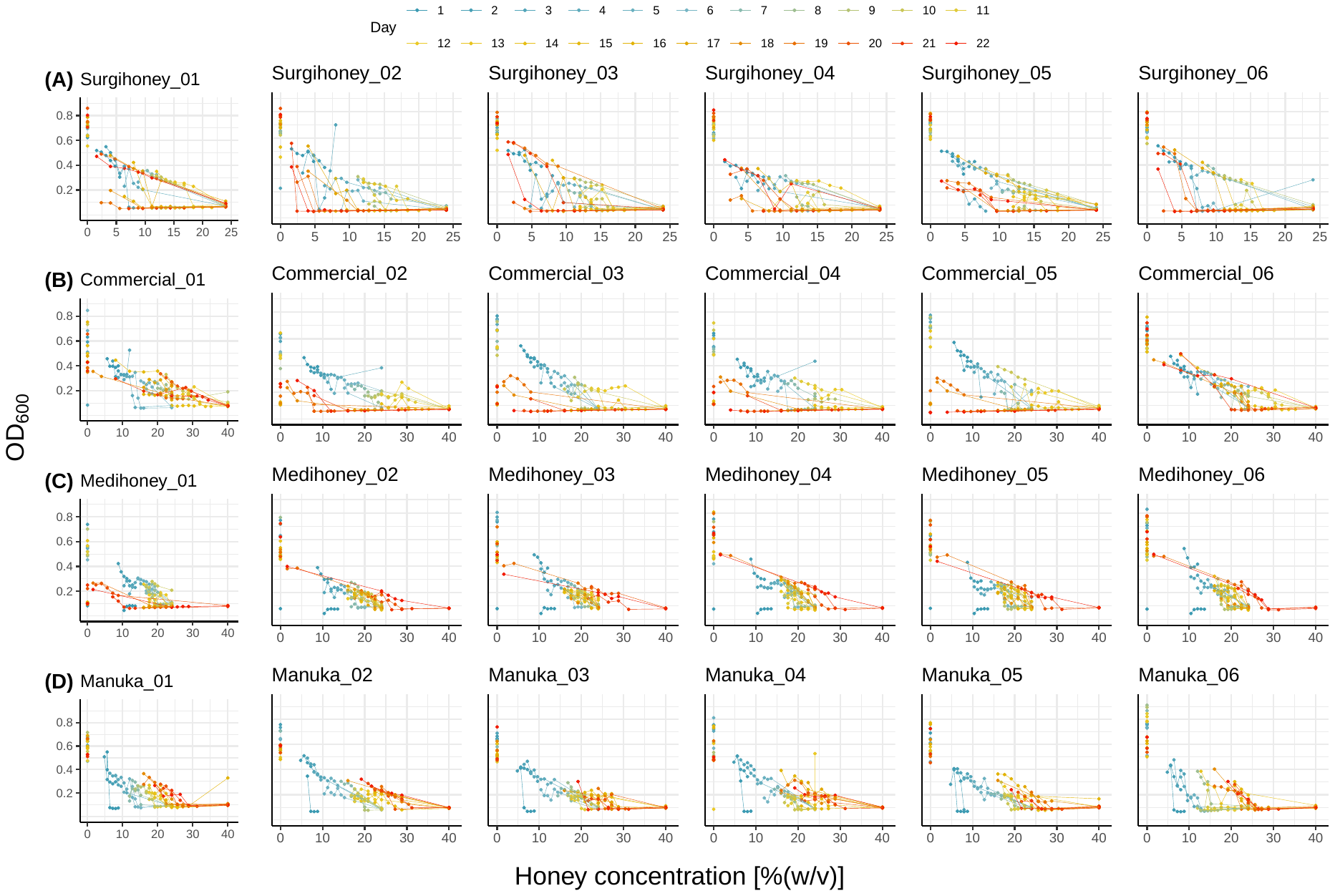

**Supplementary Figure 2.**

**Supplementary Figure 2. Population densities at different time points and honey concentrations for 24 selection lines passaged with different honeys for 22 days.** Each row of panels (A-D) shows population density, estimated by OD_600_ (*y*-axis) of six selection lines (each selection line shown in a different panel) passaged with one of the four honey compounds (each honey compound shown in a different row of panels). The *x*-axis shows the concentration of the respective honey in %(w/v), and each line gives the observed population densities across all tested concentrations on a given day (different days/lines have different colours, going from blue to green to yellow to red over earlier/later time points; shown in the legend at the top). Note that for a given selection line on a given day, there are multiple concentrations represented, because at each growth cycle there were multiple wells for each selection line (see Methods), with only the population from the well with the highest honey concentration supporting viable growth transferred to the next growth cycle (day).

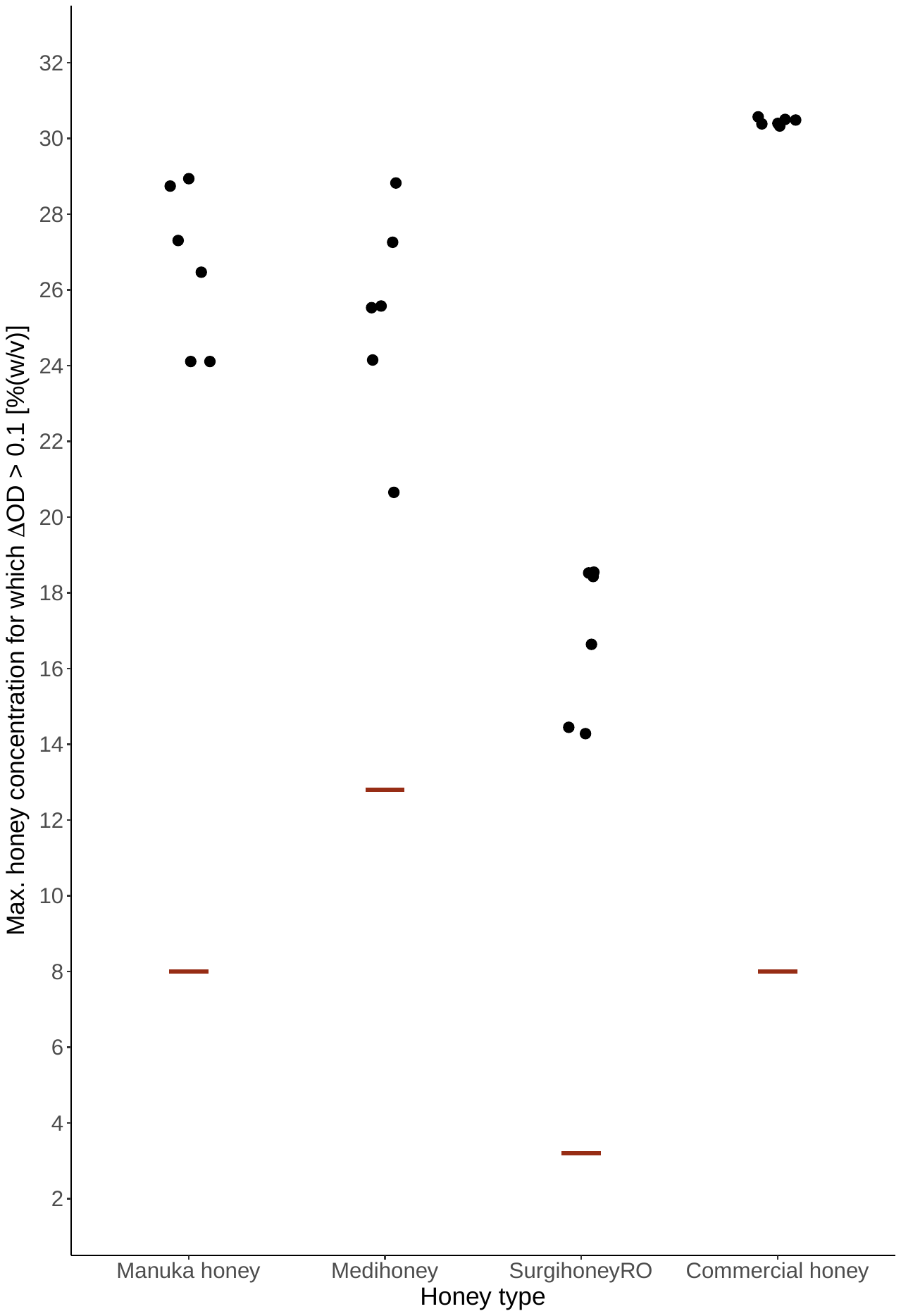
**Supplementary Figure 3.**

**Supplementary Figure 3. Highest honey concentration each selection line was able to grow at during 22d of serial passage in honey-supplemented medium.** For each honey compound, we passaged six selection lines by daily transfer into increasing concentrations of honey. Each point represents the highest concentration for which an individual selection line showed ∆OD_600_ > 0.1 after 24h incubation at any time during the experiment (points are jittered along the axes). The red line represents the initial IC_90_ of the wild type strain, *E. coli* K-12 MG1655, for each honey.

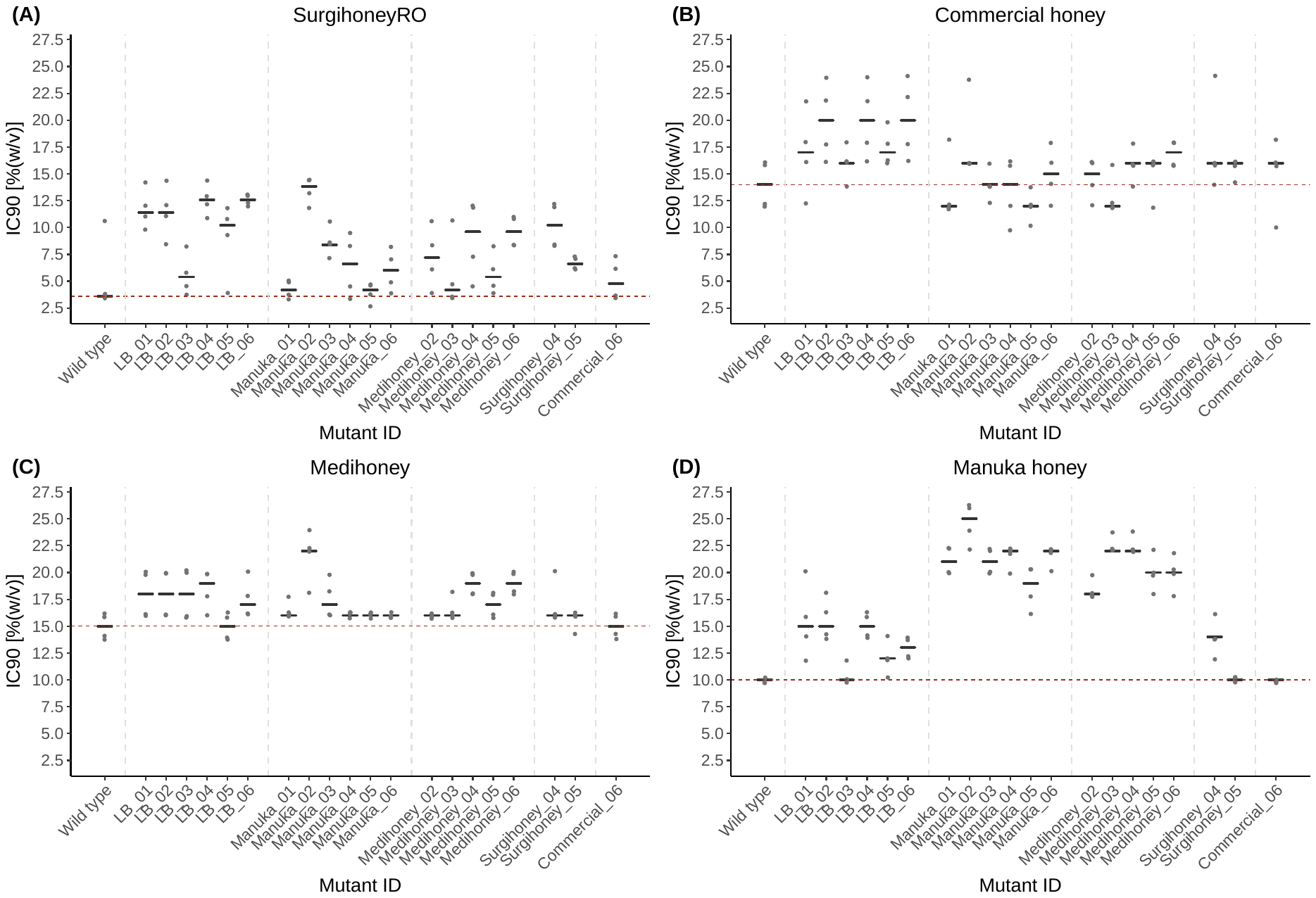
**Supplementary Figure 4.**

**Supplementary Figure 4. Resistance of serially-passaged putative resistant mutants to four different honeys.** Each of the four panels shows the IC_90_ of 14 honey-adapted (and six control-adapted) isolates and of the parental strain *E. coli* K-12 MG1655 to four honey products after 24h incubation. The black lines represent the median of four independent replicates per serially-passaged isolate, replicates are shown as dots (jittered along the *y*-axis). The red horizontal line represents the median IC_90_ of four independent replicates of the parental strain.

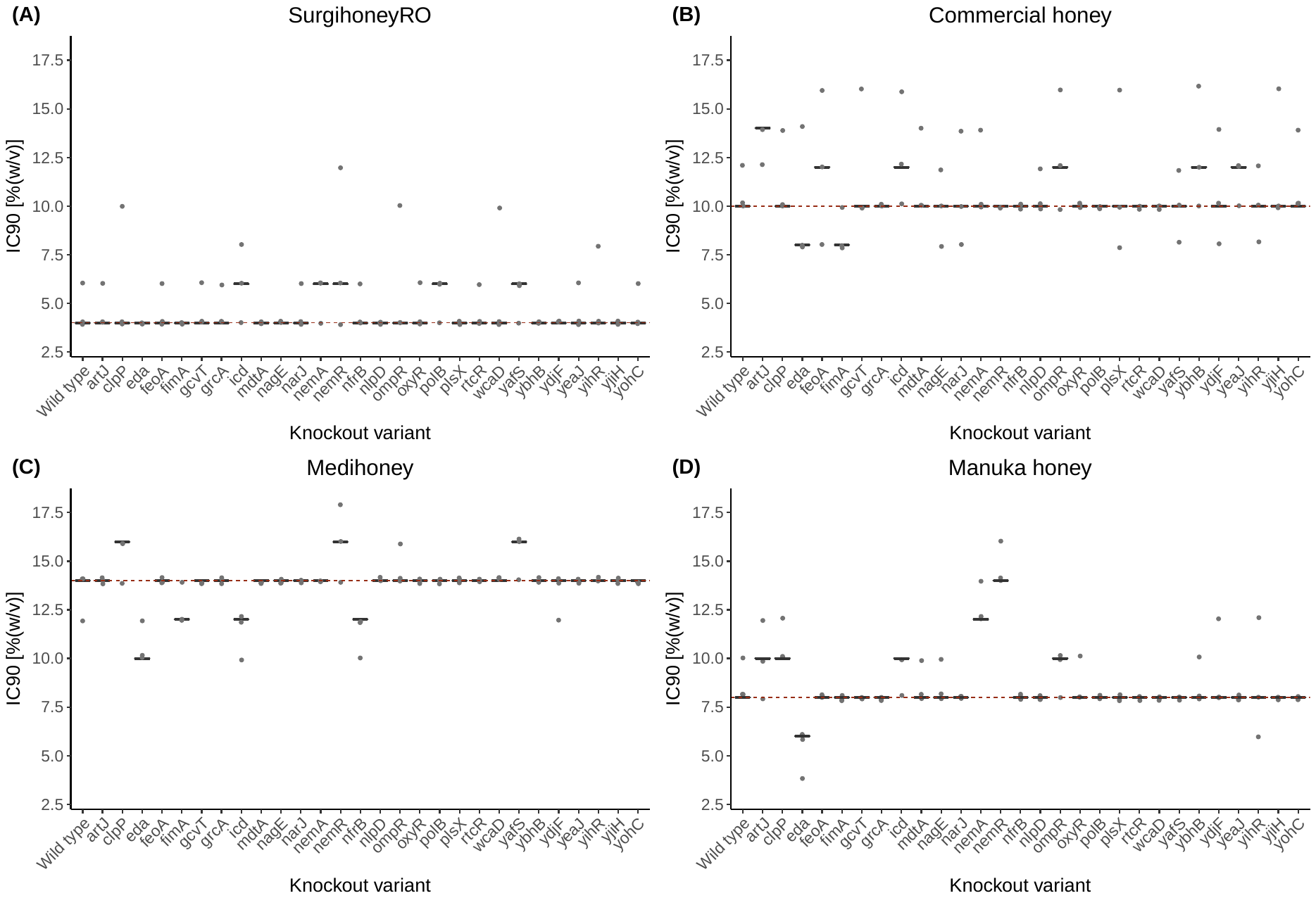
 **Supplementary Figure 5.**

**Supplementary Figure 5. Honey resistance of single-gene deletion variants.** Each panel shows the IC_90_ of 28 single-gene knockout variants from the Keio Knockout Collection and of the parental strain *E. coli* K-12 BW25113 after 24h incubation. Each black line is the median of three replicates per knockout variant; replicates are shown as dots (jittered along the *y*-axis). The dashed red line represents the median IC_90_ of three replicates of the wild-type strain, *E. coli* K-12 BW25133.

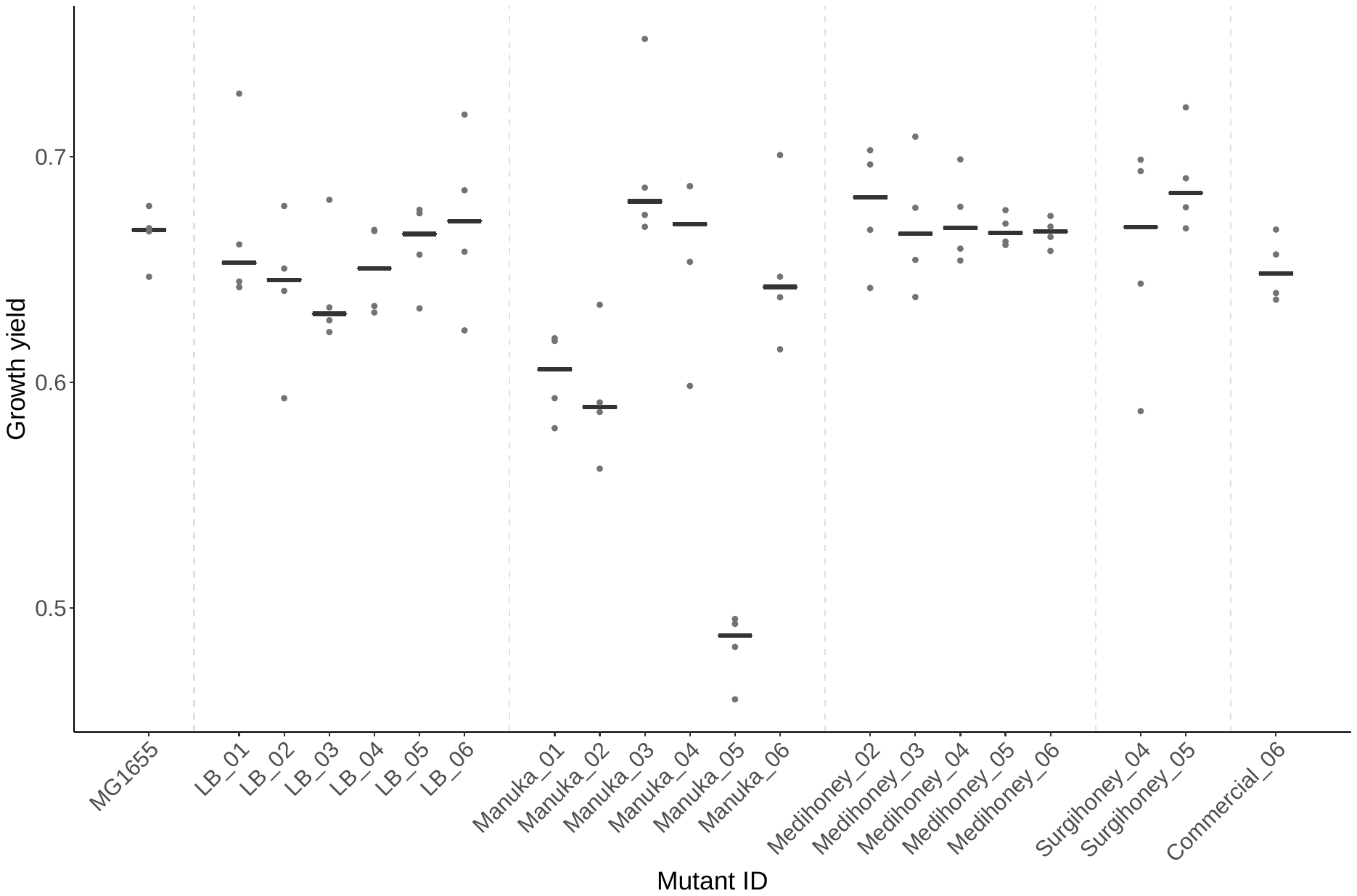
**Supplementary Figure 6.**

**Supplementary Figure 6. Growth yield of serially passaged, putative honey-resistant isolates in honey-free growth medium (LB).** Growth yield (*y*-axis) is shown for the wild type (MG1655), six isolates serially passaged in LB (LB_01-LB_06), and 14 isolates serially passaged in different honeys (labelled according to honey and replicate) (*x*-axis). Each black line represents the median of four replicates (points).

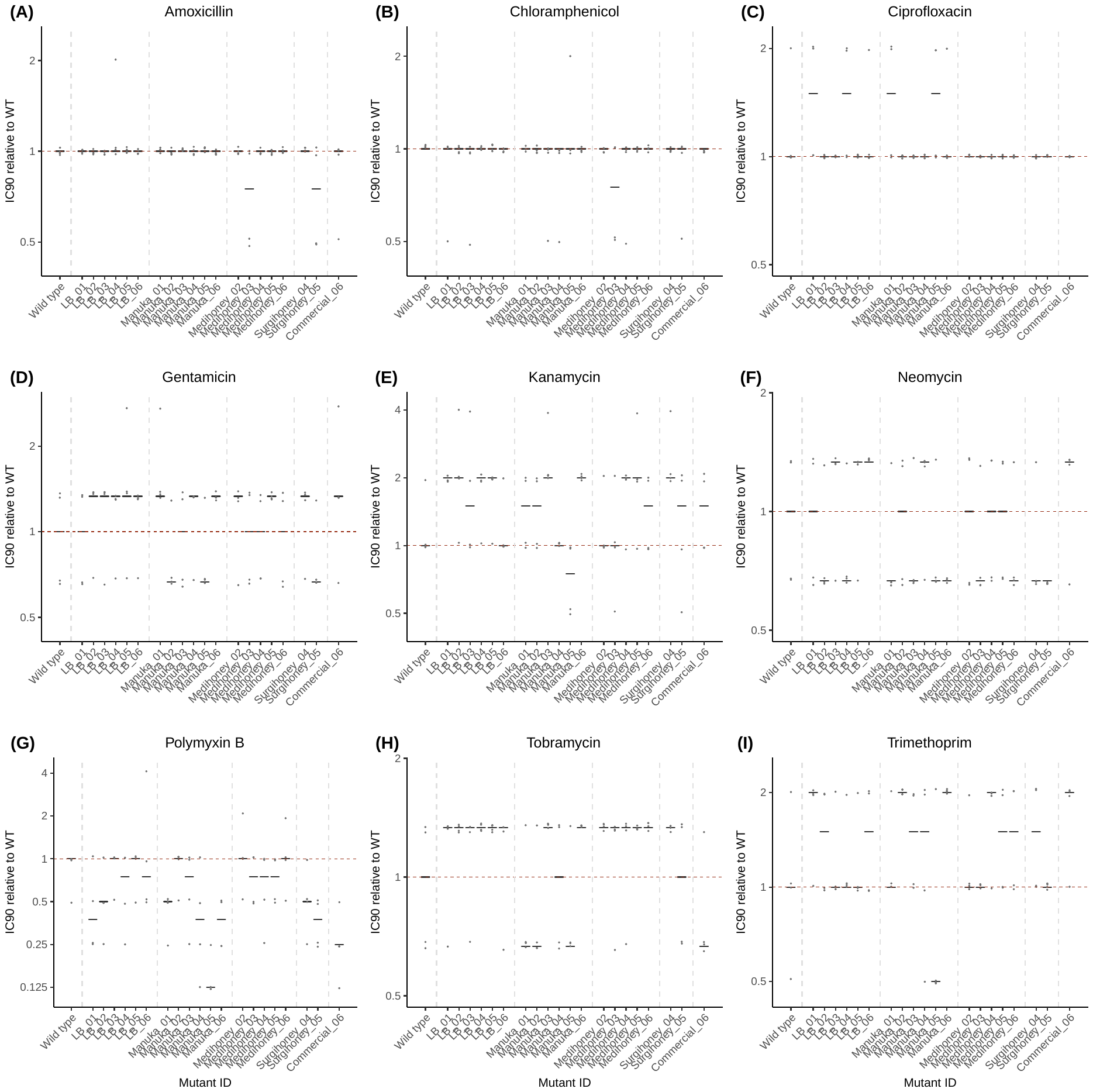
**Supplementary Figure 7.**

**Supplementary Figure 7. No evidence for cross-resistance between four honeys and nine antibiotics.** Each panel shows the IC_90_ of 20 isolates that were serially-passaged in honey or LB relative to the IC_90_ of the wild type (*E. coli* K-12 MG1655; individual replicates for the wild type are also shown relative to the median for the wild type, which is the red horizontal line in each panel) towards one antibiotic. Each black line is the median of four independent replicates per isolates; replicates are shown as dots (jittered along the *y*-axis).

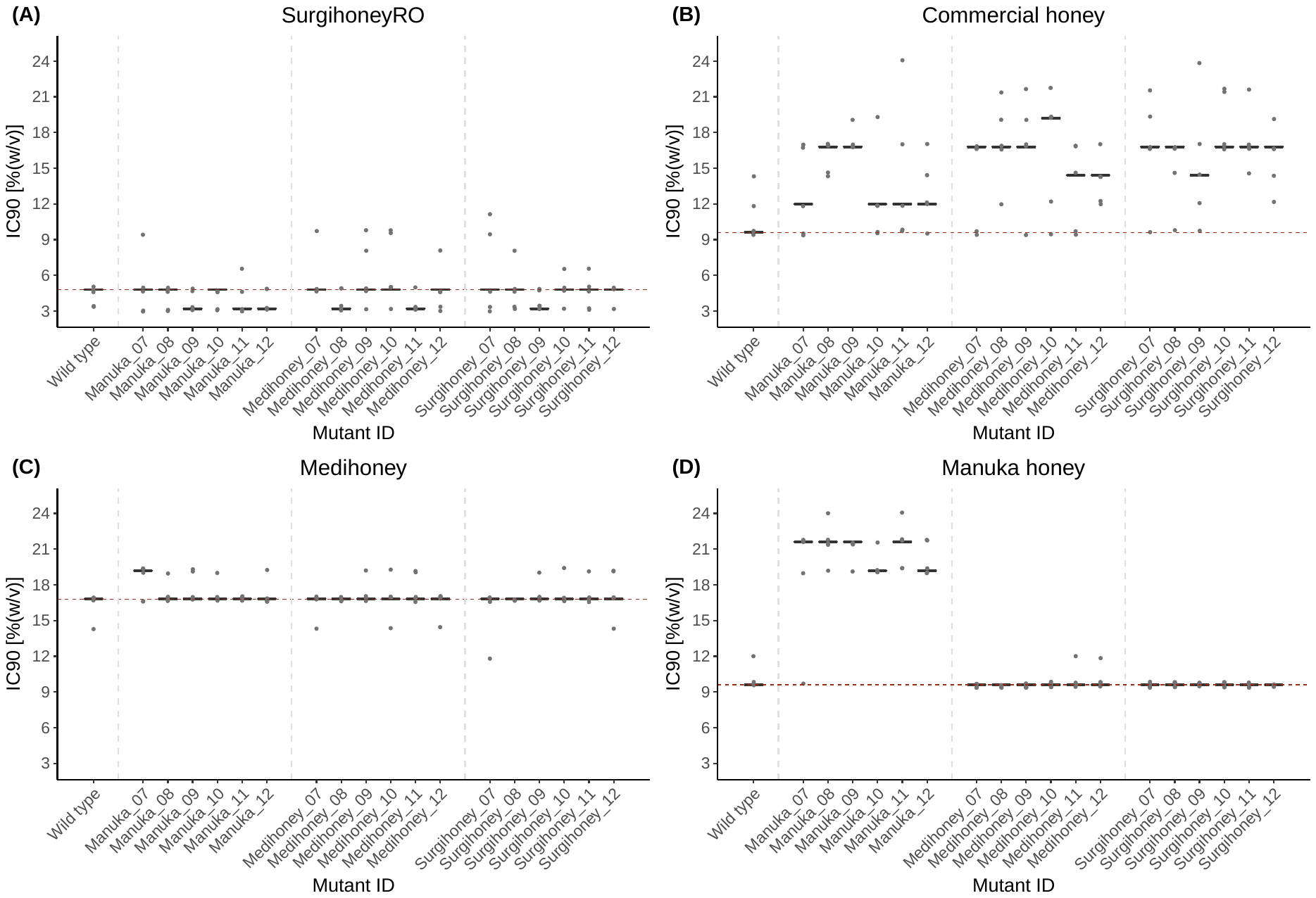

**Supplementary Figure 8.**

**Supplementary Figure 8. IC_90_ of 18 single-step putative resistant mutants to four different honeys.** Each of the four panels shows the IC_90_ values of 18 single-step mutants and of the parental strain *E. coli* K-12 MG1655 to four different honeys. The black lines represent the median of five independent replicates per single-step putative resistant mutant, replicates are shown as dots (jittered along the *y*-axis). The red horizontal line represents the median IC_90_ of five independent replicates of the wild-type strain.
